# Traversing the canopy: phenology-driven changes and within-canopy transport shape the phyllosphere microbiome in a temperate floodplain hardwood forest

**DOI:** 10.64898/2026.03.26.714518

**Authors:** Dinesh S. L. Chetti, Beate Michalzik, Marina M. Sandoval, Patrick Zerhusen, Ronny Richter, Rolf A. Engelmann, Tom Künne, Christian Wirth, Kirsten Küsel, Martina Herrmann

## Abstract

- Phyllosphere microbiomes are subject to microbial import from various sources and undergo substantial changes during phenological changes of plants. However, these processes are still poorly understood for forest canopies. We propose that phenology-driven changes in host properties, and rainwater-mediated, within-canopy transport shape the phyllosphere microbiome in temperate forests. Leaves and throughfall samples were collected from oak, ash and linden trees at top, mid, and bottom canopy positions at the Leipzig canopy crane facility (Germany) at time points representing early, mid and late phenological stages. Bacterial community composition was assessed by 16S rRNA gene amplicon sequencing.
- Phenological stages explained 19% of phyllosphere bacterial community variation, followed by tree species identity (12%) and canopy position (2%). Later phenological stages exhibited more homogeneous and functionally redundant phyllosphere communities along with a strong decline of plant pathogens and increasing potential for microbially mediated biocontrol mechanisms. Throughfall transported up to 10^11^ bacterial cells per litre with maximum bacterial fluxes at the canopy top.
- Our findings demonstrate that in temperate forests, phenology-driven effects on the phyllosphere microbiome are far more important than tree species’ specific effects. Extent and selectivity of throughfall-mediated mobilization may play a crucial role for the spatial heterogeneity of microbial communities in tree crowns.

## Introduction

Tree canopies represent the interface to the atmosphere and are exposed to the input of energy, water and matter, spanning a surface area of approximately 4 x 10^8^ km^2^ on a global scale (Morris and Kinkel, 2002). Trees harbour habitats on leaves (phyllosphere) and bark, which are host to microbial communities involved in central biogeochemical processes as well as plant-microbe interactions, affecting plant community dynamics, ecosystem functioning and productivity (Laforest-Lapointe and Whitaker, 2019). Bacteria present in the phyllosphere interact with each other, the plant host, and the environment through competition, commensalism, antagonism and neutral interactions (Schäfer et al., 2022), which includes various functions beneficial to plants such as nitrogen fixation (Carrell and Frank, 2014), methane oxidation (Iguchi et al., 2015), and phosphate solubilization (Venkatachalam et al., 2016), but also pathogenesis, decreasing the host’s fitness (De Mandal and Jeon, 2023).

Tree species identity, canopy position, and phenology have been reported as strong drivers for phyllosphere bacterial communities in temperate forests (Al Ashhab et al., 2021; Ginnan et al., 2022; Herrmann et al., 2021; Laforest-Lapointe et al., 2016). However, the relative importance of these controls is still poorly understood. Technical challenges in accessing the whole tree crown for sampling have so far largely constrained our understanding of vertical patterns of the phyllosphere microbiome (Stone and Jackson 2019, Herrmann et al. 2021). Especially the canopy top represents a key interface between the forest and the surrounding atmospheric environments, mediating microbial and chemical exchange while its phyllosphere microbiome is the most exposed to harsh weather conditions and UV radiation.

Host physiology and leaf chemistry are central factors underlying the influence of tree species identity on phyllosphere bacterial communities. These host-specific properties are subject to phenology-driven changes, e. g., reflected by an increased release of sugars, amino acids, or inorganic ions from leaves towards leaf senescence (Tukey, 1970; Van Stan et al., 2012), which in turn affects growth and competition of phyllosphere-associated microorganisms and thus their successional dynamics (Feeny, 1970; Mediavilla and Escudero, 2003; Sancho-Knapik et al., 2023). These processes are central to phyllosphere community assembly, which is typically dominated by stochastic processes (Chen et al., 2025; Deng et al., 2024), including the limitation of bacterial movement through physical, biological, or environmental constraints (dispersal limitation; (Li and Gao, 2023)), or random changes in bacterial community composition due to replication, lysis and colonization (drift) (Chase and Myers, 2011).

The phyllosphere receives bacteria from various sources. These include atmospheric sources through air–borne inputs such as including precipitation and dust particles (Ponette-González et al., 2020; Zhou et al., 2021), while herbivorous and pollinator insects constitute another import vector (Feeny, 1970). In tree canopies, incoming rainfall is partitioned into throughfall, stemflow and interception loss (Levia et al., 2011). Throughfall is rich in nutrients (C, N and P) (Lusk, 2024) and may harbour up to 100 times more bacteria than rainwater (Bittar et al., 2018). Consequently, throughfall is likely to act as central vector in the translocation of microorganisms, and translocation may be taxon-specific, as the ability of bacteria to remain attached or be washed–off from leaf surfaces depends on their ability to form surfactants (Hassan and Frank, 2003) or biofilms (Lyautey et al., 2005; Pantanella et al., 2006). Yet, these translocation processes have only rarely been studied, and observations were typically confined to collecting throughfall and the transported microbial communities below the canopy (Krüger et al., 2025; Stone and Jackson, 2019; Teachey et al., 2018), lacking insight into the relevance of throughfall for within canopy distribution processes.

Here, we propose that phenology-driven changes in plant host properties, and transport of microorganisms within the canopy via throughfall are key mechanisms driving canopy microbial diversity and its spatio-temporal variation in a temperate forest. We leveraged the infrastructure of the Leipzig canopy crane facility, installed in a floodplain hardwood forest, which allows full access to tree canopies of up to 35 m height, and we employed a novel approach of throughfall sampling within tree crowns at different height levels. Combined with sampling of the phyllosphere, this approach enabled us to assess vertical patterns of phyllosphere communities and their potential connection by throughfall-mediated transport processes. Focusing on three common tree species, *Quercus robur* L. (English oak), *Fraxinus excelsior* (European ash) and *Tilia cordata* MILL (Winter linden), we followed bacterial communities in phyllosphere versus throughfall across early (May), mid (July) and late (October) phenological stages at three different height levels within the canopy - top, mid, and bottom. We aimed to (i) disentangle the controls of canopy position and tree species identity on the phyllosphere microbiome across different phenological stages, (ii) follow introduction and succession of bacterial taxa and quantify the relative importance of different community assembly mechanisms, and (iii) evaluate the contribution of throughfall-mediated bacterial transport to phyllosphere microbiome patterns.

## Methods

### Site description

The sampling site is located in Burgaue, Northwestern part of Leipzig (Saxony, Germany) on the floodplains of the Elster, Pleisse, and Luppe rivers. The sampling site experiences a continental climate with the mean annual temperature of 8.4°C and mean annual precipitation of 516 mm (Jansen, 1999). Annual precipitation in 2021 was 539 mm (Richter et al., 2022), making it slightly wetter than the long-term average. The Canopy Crane site consists of 18 different tree species and around 800 tree individuals in an area spanning 1.65 km^2^ (Patzak et al., 2020). The most abundant tree species in the forest are European ash (*Fraxinus excelsior* L.) and English oak (*Quercus robur* L.) (Richter et al., 2016), which are also the focal trees of this study along with *Tilia cordata* MILL (Winter linden). Hereafter, the tree common names are simply referred to as oak, ash and linden throughout the manuscript. The study area represents a mature old-growth forest with diameter at breast height (DBH) values of the sampled individuals ranging from 55 to 136 cm for oak, 68.7 to 84 cm for ash and 71 to 81 cm for linden. (Table S1).

### Sample collection and processing

Leaf samples were collected from each three individuals of oak, ash and linden trees, at three different time points during the vegetation period, representing an early phenological stage directly after leaf sprouting (May), a mid-phenological stage in July and a late phenological stage in October 2021 with clear signs of senescence on the leaves (Figure 1). In addition, to follow potential vertical changes in phyllosphere microbiome composition across the tree crowns, we sampled the top, middle and bottom canopy positions with each three spatial replicates. The different canopy positions were accessed using a gondola operated by a crane with a maximum sampling height of about 35 m. At each sampling position, approximately ten leaves were randomly collected from different twigs which were approximately 50 cm apart from each other using sterile scissors and were transferred to autoclaved polypropylene bottles. Throughfall samples were collected at the same time points as leaf samples and additionally in March 2021 (hereafter referred as March–throughfall). Throughfall collectors (sterile polypropylene bottles) with a funnel size of 113 cm^2^ surface area were deployed close to the locations of phyllosphere sampling (top, middle and bottom) within the canopy and exposed for a period of 14 days until throughfall samples were collected. Samples of rainwater which had not been in contact with the tree crowns were collected above the forest canopy by placing sterile polypropylene bottles on top of the crane (referred as “crane top”). Our sampling design allowed the collection of both dry deposition and rainwater-associated transport of microorganisms as well as the integrated sampling of multiple rain events during the exposure period. As sampling directly after single rain events was not possible with the infrastructure used, we cannot entirely rule out that growth of microorganisms in the throughfall contains may have led to a slight overestimation of throughfall bacterial abundances. The placement height of the throughfall collectors depended on the canopy architecture and tree individuals and varied from 26–34 m for the top position, 22.5–29.5 m for the mid position, and 19.5–24 m for the bottom position (File S1). Both leaf and throughfall samples were transported in a cooled condition with ice packs, and stored at 4 °C upon arrival to the lab.

**Figure 1.**
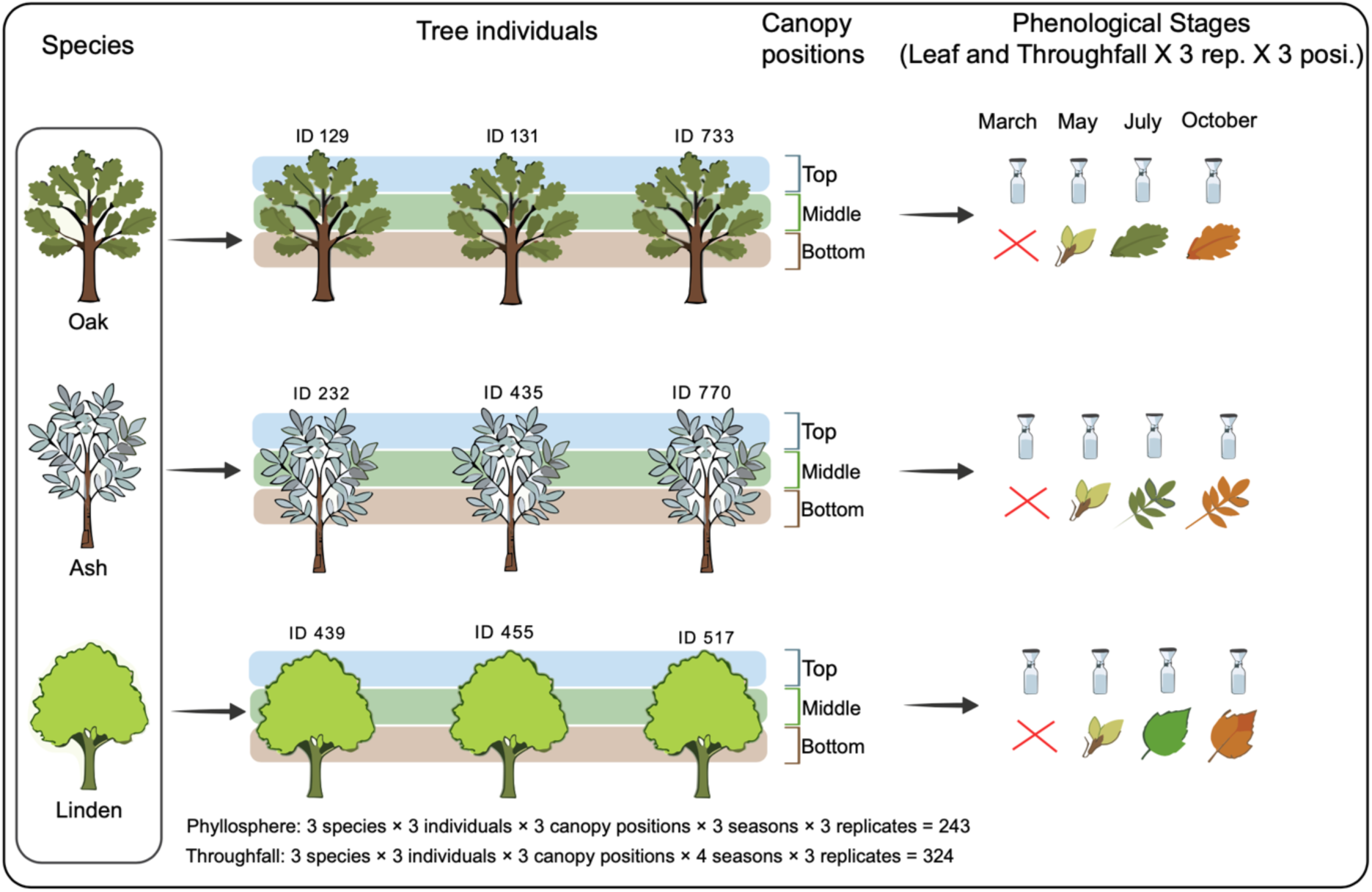
Study design and sampling strategy. Leaf and throughfall samples were collected from three individuals of oak, ash and linden trees at three canopy positions (Top, middle and Bottom) in three replicates per position during May, July and October, representing early, mid and late phenological stages respectively. Throughfall samplers were installed at the respective canopy positions 14 days prior to the day of sampling. Additional throughfall samples were also collected in March, when the trees were leafless.

### Sample processing, DNA extraction and amplicon sequencing

For detachment of leaf-associated epiphytic bacteria, leaf samples were immersed in 250 ml of suspension buffer (0.15 M NaCl, 0.1% Tween 20) in the collection bottles. These bottles were subjected to brief sonication twice for 1 min at 10% intensity, followed by shaking for 20 min at 100 rpm at room temperature to enhance the detachment of bacterial cells from the surface of the leaves. The resulting suspensions were filtered through 0.22 µm pore size PES (polyethersulfone) membrane filters (Supor, Pall Corporation) to collect the bacterial cells. The remaining leaves were used for determination of dry weight after drying at 50 °C for one week. Throughfall samples were directly filtered through the same filter type. Filters were stored at −80 °C until DNA extraction.

DNA was extracted from the filters using the DNeasy PowerSoil Pro Extraction kit (Qiagen) following the manufacturer’s protocol. We used the primers Bakt_0341F (Klindworth et al., 2013) and Bakt_0799R (Chelius and Triplett, 2001; Herrmann et al., 2021) for the amplification of the V3–V4 of bacterial 16S rRNA genes, with the reverse primer being designed in order to avoid the amplification of chloroplast-related 16S rRNA genes. The PCR amplification, paired–end library preparation and amplicon sequencing (2×300 base pairs; v3 chemistry) were carried out using an Illumina MiSeq sequencer based on a protocol described by Krüger et al., 2021. The demultiplexed sequence files from this study were deposited in the European Nucleotide Archive (ENA) (accession number: PRJEB109949 and PRJEB109950). Quantitative PCR was performed using the primer combination Bakt_0341F (Klindworth et al., 2013) and Bakt_0799R (Chelius and Triplett, 2001) under the following cycling conditions, 95 °C for 10 minutes, followed by 45 cycles of 95 °C for 30 s, 53 °C for 30 s, and 72 °C for 40 s, followed by melting curve analysis. qPCR was performed using Brilliant SYBR Green II Master mix (Agilent Technologies) on a Bio–Rad CFX96 platform (BioRad).

### Sequence analyses

We analysed raw sequence reads from rainwater (“crane top”), phyllosphere and throughfall samples using the Divisive Amplicon Denoising Algorithm 2 v1.26 dada2 R package (Callahan et al., 2016). We performed primer removal (*truncLeft(17,21*)), trimming (*filterAndTrim*; truncLen=c(280, 220)), error learning (*learnErrors*, *maxEE(2,4)*), denoising (*dada*), amplicon sequence variant (ASV) inference (*dada*), merging (*mergePairs*), table generation (*makeSequenceTable*), removal of chimeric reads (*removeBimeraDenovo*) and classification (*assignTaxonomy* using silva_nr99_v138.1 and silva_species_assignment_v138.1 databases) using dada2. The final count table for the crane top, phyllosphere and throughfall consisted of 10, 196 and 319 samples, respectively. Crane top, phyllosphere and throughfall count tables were rarefied to 19780, 4217 and 4441 reads per sample, respectively based on the sample with lowest number of sequence counts. The count tables were further used for cell abundance estimations based on the following calculation, [relative abundance/average 16S rRNA operon number (at family level)] X 16S rRNA gene abundance data determined by qPCR. Briefly, the relative abundance values were divided by the family level mean 16S rRNA operon numbers (https://rrndb.umms.med.umich.edu/), then multiplied with qPCR abundance data. The ASVs were categorised into abundant (> 0.1 %), intermediate (0.01 to 0.1 %) and rare (< 0.01 %) (Jiao and Lu, 2020) based on their relative abundance in the phyllosphere communities. ASVs that repeatedly occurred in consecutive phenological stages were referred as ‘successional’ ASVs, while the unique ASVs were referred to as ‘newly introduced’ ASVs. The number of common and unique ASV present across phenological stages (early–mid and mid–late phenological stages) and vertically (top–mid and mid–bottom canopy positions) were assessed to understand the phenological and throughfall mediated vertical transfer associated changes, respectively.

### Functional prediction using PICRUSt2 analysis

Estimated cell abundances of bacterial ASVs, calculated as described above, were used for functional prediction using PICRUSt2 (Phylogenetic Investigation of Communities by Reconstruction of Unobserved States) (Douglas et al., 2020). In brief, the Kyoto Encyclopedia of Genes and Genomes (KEGG) ortholog (KO) gene families were predicted through the picrust2_pipeline.py function with the option of skipping operon normalisation (--skip-norm), since the input data was already copy number normalized. The assignment of KO families to various KEGG levels was carried out using the mapping file from PICRUSt2 Github (accessed on 14.05.2025). Pathway abundances (KEGG level 3) were compared between tree species and across phenological stages and tested for significance using Kruskal–Wallis and Dunn’s pairwise test.

### Community assembly process analysis

The assembly process analysis was carried out based on absolute abundance data using the phylogenetic bin–based null model analysis (iCAMP) R package (Ning et al., 2020), which was developed based on a previous study (Stegen et al., 2013) to quantify the relative importance of ecological processes (deterministic: homogeneous selection (HoS) and heterogeneous selection (HeS), stochastic: dispersal limitation (DL), homogenizing dispersal (HD) and drift (DR)) governing phyllosphere microbiome community assembly across the phenological stages. Absolute abundance data were divided by a factor of 23.5 prior to analysis to avoid computational issues associated with large numbers. Briefly, phyllosphere ASVs were clustered into bins based on their phylogenetic similarities. These bins were subjected to an abundance–based null model analysis to calculate the beta net relatedness index and Raup–Crick through 999 pairwise iterations to infer the relative importance of deterministic (HoS: βNTI < −1.96, and HeS: βNTI > 1.96) and stochastic (|βNTI| < 1.96) (HD: RC < −0.96; DL: RC > 0.96; DR: |RC| < 0.96) ecological processes governing bacterial community assembly in the phyllosphere.

### Statistical analyses and visualization

Principal coordinate analysis (PCoA) was performed using the vegan package (Oksanen et al., 2020) in R to analyze the clustering of samples in the ordination space across phenological stages. Singletons were removed before performing PCoA analysis. Permutational ANOVA (PERMANOVA) analysis (999 iterations) was performed to assess the contribution of phenological stages, tree species identity, and canopy position to the total variation in bacterial community structure using vegan. Phenological stage-wise PERMANOVA tests were also conducted to assess the relative contribution of tree species and canopy position to bacterial community variability for each month separately. Normality test (Shapiro-Wilk test, *p*<0.05) was performed before selecting one way ANOVA Tukey’s HSD pairwise test/Kruskal–Wallis and Dunn’s pairwise test for analysing total abundance, family level abundance and throughfall bacterial flux changes across the phenological stages and canopy positions. We generated boxplots using ggplot2 and performed Kruskal–Wallis and Dunn’s pairwise test to identify significant changes in bacterial diversity (species richness and Shannon’s diversity H index) across phenological stages. Venn diagrams were used to understand the percentage of successional and unique ASVs across phenological stages using eulerr R package. The bacterial abundance changes across the phenological stages were tested using Kruskal–Wallis and Dunn’s pairwise test. Heatmaps with hierarchical clustering for ASVs (based on Euclidean distances) were created using the pheatmap R package to follow abundance changes (log transformed) across the phenological stages. Family level abundance changes across canopy positions and phenological stages were tested through Kruskal–Wallis and Dunn’s pairwise test using the package stats. Key ASVs which increased or decreased in terms of abundance across the phenological stages were identified based on linear discriminant score (LDA) using linear discriminant analysis effect size (LEfSe, microbiomeanalyst). Key ASVs that preferentially occurred in crane top samples (rainwater), in the phyllosphere or in throughfall were analysed by comparing the relative abundances of common ASVs between any two selected groups (crane top vs throughfall at the top canopy position or phyllosphere vs throughfall at all three canopy positions) using LEfSe analysis. The throughfall bacterial flux was calculated as per sampling event by factoring the estimated cell abundances per L, rainwater/throughfall volume during the sampling period, and surface area of the collectors. The KEGG assignment resulting from PICRUSt2 analysis was carried out at category (level 1), sub–category (level 2) and pathway (level 3) levels. KEGG pathway abundance changes at level 3 were analyzed pairwise (early–mid and mid–late phenological stages) for each tree species using Kruskal–Wallis and Dunn’s pairwise test, *P*<0.05 using stats R package. The KEGG pathway abundance (log_10_–transformed) change across phenological stages was visualised using pheatmap package in R.

## Results

### Changes in phyllosphere bacterial community patterns and abundance across early, mid, and late phenological stages

Across all three tree species, phenological stage was the most important factor driving phyllosphere bacterial community composition, as it explained 19.23% of community variability, followed by tree species identity (12.18%). In contrast, canopy position (2.34%) and tree individual (1.15%) constituted only minor contributions to community variability (Figure 2A, Table S2). Phyllosphere communities sampled in May, July, and October were clearly distinct with a slightly higher similarity between bacterial communities of the mid and late phenological stages compared to the early phenological stage. When analyzing community drivers separately for each phenological stage, tree species identity explained most of the bacterial community composition variability with a gradually increasing effect from the early (19.48%) to mid (23.66%) and late (25.88%) phenological stage (Figure 2B, Table S2). The contribution of canopy position was similar across the phenological stages and ranged from 4.99% (May) to 5.86% (July) (Table S2).

**Figure 2.**
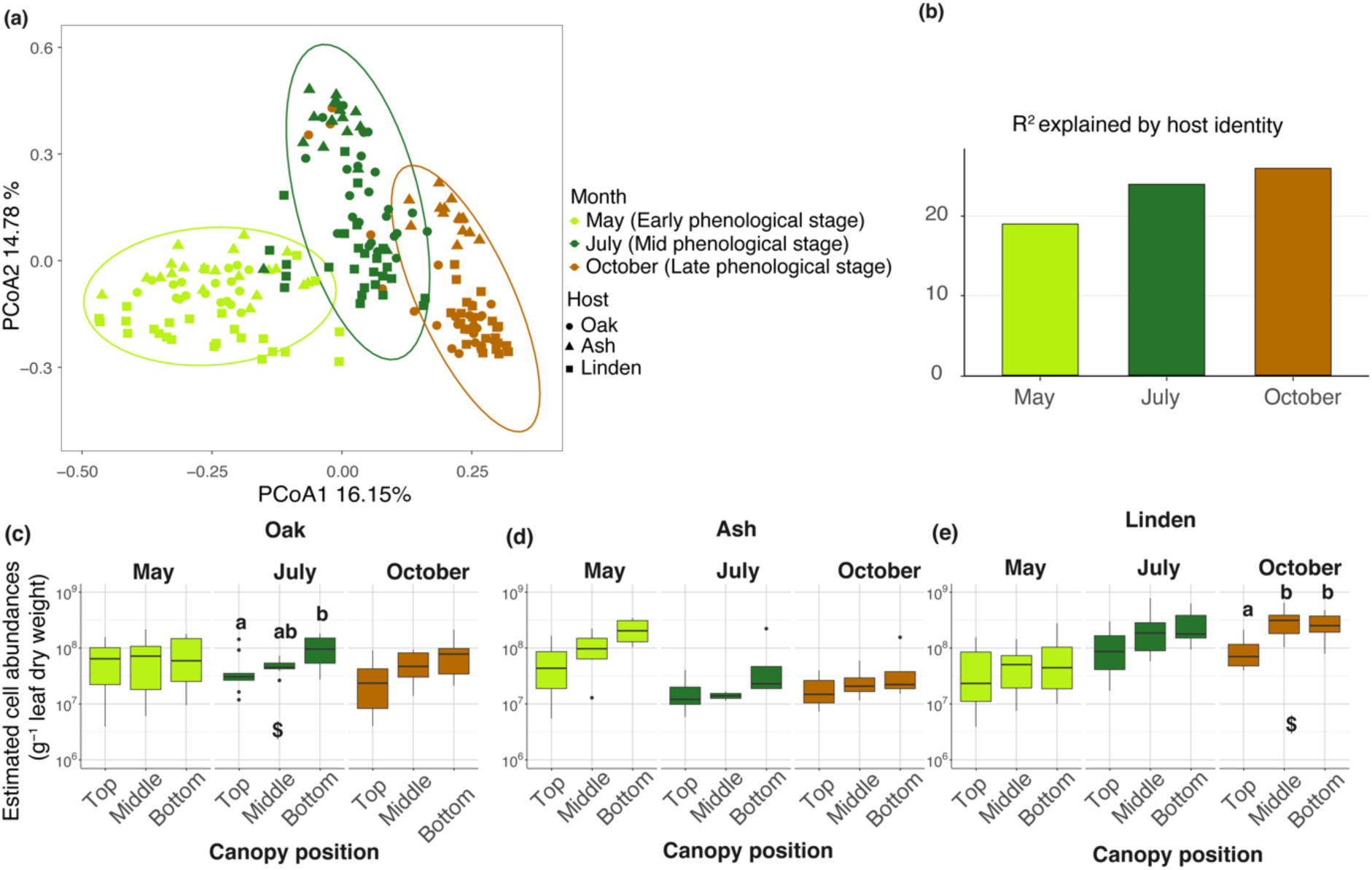
Principal co-ordinate analysis of the phyllosphere bacterial communities associated with oak (*Q. robur* L.), ash (*F. excelsior*) and linden (*T. cordata*) across different phenological stages. (a) Sample-based ordination plot for phyllosphere bacteria (operon normalised) generated from amplicon sequence variants (ASV) compositions. Ellipses represent 95% confidence intervals for the tested factor variable (i.e. Phenological stages). (b) Community variation (R^2^) (0–100) explained based on host identity according to phenological stages, May (Early), July (Mid) and October (Late phenological stages). (c-e) Estimated cell abundances of phyllosphere bacteria across tree species and canopy positions. Alphabets denote statistical difference between canopy positions. ^$^ depicts statistical significance (*P*<0.05) based on Kruskal’s wallis and Dunn’s pairwise test (*P*<0.05).

Despite the rather small effect of canopy position on phyllosphere community composition, canopy position significantly influenced Shannon diversity (H’) with an increasing trend from the top to the bottom canopy position of the oak and linden across all phenological stages, while for ash, phyllosphere community diversity was highest at the middle canopy position during mid and late phenological stages (Kruskal–Wallis and Dunn’s pairwise test, *P*<0.05) (Figure S1). Similarly, estimated abundances of phyllosphere bacteria were significantly higher at the bottom compared to the top canopy position during the mid and late phenological stages for oak and linden (Kruskal–Wallis and Dunn’s pairwise test, *P*<0.05) while for ash, such a vertical trend was confined to the early phenological stage (Figure 2C, Figure 2E and File S2).

Following changes in Shannon’s diversity index H’ and estimated abundances across phenological stages revealed that Shannon diversity was significantly higher during the mid (oak, ash, and linden) and late phenological stages (oak, linden) compared to the early phenological stage (Figure S1). In contrast, we observed opposing trends of phyllosphere bacterial abundances over time. For ash, the bacterial abundances in the phyllosphere decreased during mid and late phenological stages compared to the early phenological stage, while the opposite pattern was observed for linden (Kruskal–Wallis and Dunn’s pairwise test, *P<*0.05), and bacterial abundances in the oak phyllosphere remained on a similar level across all phenological stages (Figure 2C–2E).

### Compositional changes and successional transfer across phenological stages

Bacteria belonging to 20 phyla, 203 families, and 527 genera were detected in the phyllosphere samples of this study. Ninety–five percent of the total reads belonged to the phyla Proteobacteria (68%), Bacteroidota (14%) and Actinobacteriota (13%), with *Sphingomonadaceae*, *Beijerinckiaceae*, *Pseudomonadaceae*, *Erwiniaceace*, *Oxalobacteraceae*, *Acetobacteraceae, Microbacteriaceae* and *Kineosporiaceae* representing the most abundant families. The relative abundance of top 20 bacterial families ranged from 74.6 to 96.3% of estimated cell abundances g^-1^ of leaf dry weight (Figure S2), with substantial variations across the phenological stages and tree species. The estimated cell abundances of bacteria affiliated with *Beijerinckiaceae* showed an increase during the mid and late phenological stages across all tree species (Figure S3), being highest at the bottom canopy position for specific phenological stages and tree species (Kruskal–Wallis and Dunn’s pairwise test, *P*<0.05 and ANOVA test, *P<*0.05) (Table S3). In contrast, the abundance of *Erwiniaceae* decreased during the mid and late phenological stages but without any consistent trends in the vertical distribution (Figure S3).

To assess the successional transfer of ASVs across the phenological stages, we analyzed the shared ASVs of the phyllosphere communities between May and July or July and October. The fraction of shared ASVs was typically higher between the mid and late phenological stage (22–43%) compared to the transition from the early to the mid phenological stage (16–29%) at all canopy positions and across all three tree species (Figure 3A). The cumulative relative abundance of successional ASVs, defined as ASVs that repeatedly occurred in consecutive phenological stages, was higher (50–92%) than that of newly introduced ASVs (8–50%) for both the July and the October communities (Figure S4) and Table S4). In turn, the relative abundance of newly introduced ASVs was higher for the mid (16–50%) compared to the late (8–13%) phenological stage (Figure S4). The successional ASVs associated with oak and linden mostly belonged to the category of abundant taxa (>0.1%) (23–49% of all ASVs), while for ash, successional ASVs were mostly contributed by ASVs of the category of rare (<0.01%) or intermediate taxa (<0.1 % and >0.01%; 39–78% of all ASVs) (Figure S5 A and Table S4). Newly introduced ASVs mostly belonged to the category of rare ASVs (<0.01%) (35–91%), irrespective of phenological stage or tree species (Figure S5 B).

**Figure 3.**
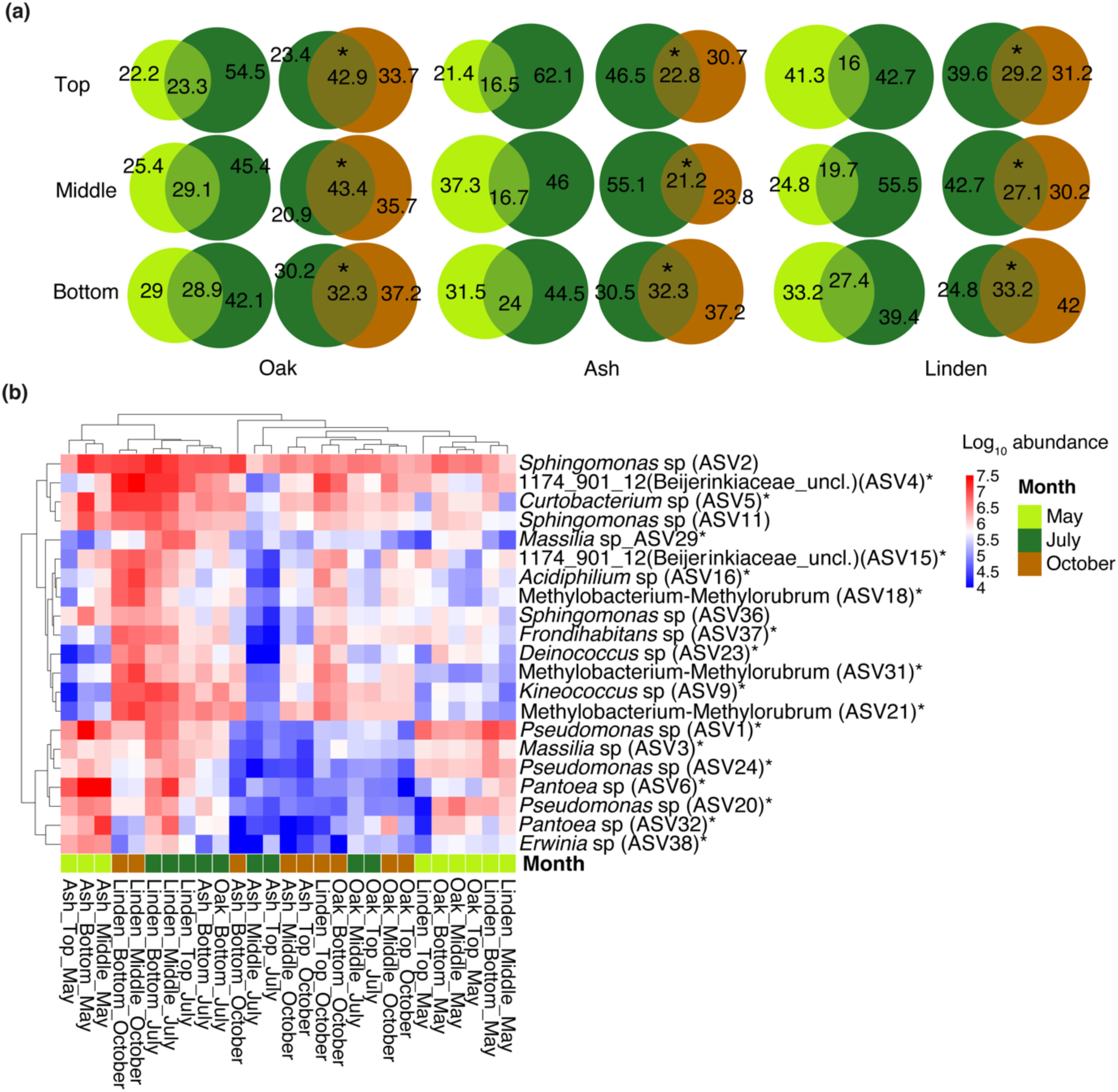
Top and shared amplicon sequence variants across phenological stages in the phyllosphere. (a) Venn diagram showing the shared and unique amplicon sequence variant (ASV) percentages among May, July and October. “Top”, “Middle” and “Bottom” denote canopy positions. May, July and October indicate phenological stages. * indicates statistical significance of shared ASV percentages between May-July and July-October based on two-proportion z-test (*P*<0.05). (b) heatmap showing the top phyllosphere amplicon sequence variants (ASVs). The colour intensity of the palette indicates the log_10_ estimated cell abundances from low (blue) to high (red). * denotes differentially abundant (*P*<0.05) phyllosphere ASVs based on LEfSe analysis.

The heatmap of the distribution patterns of the 20 most abundant ASVs showed a more distinct clustering of samples from the mid to late compared to the early phenological stage, mirroring the observations of the shared ASV (%) trends (Figure 3B). When moving from early to late phenological stages, abundance of unclassified *Beijerinckiaceae* increased, while abundances of potential plant pathogens, such as *Erwinia* sp. (ASV38), *Pantoea* sp. (ASV32) and *Serratia* sp. (ASV33) decreased (Figure 3B and File S3) (LEfSe, *P*<0.05). Specific abundant ASVs were newly introduced only during mid and late phenological stages across tree species, or appeared specific to oak, ash, or linden (Table S5).

### Drift dominates the community assembly processes in the tree phyllosphere during mid and late phenological stages

Community assembly process analysis was performed to understand the relative importance of deterministic (homogenous selection and heterogenous selection) and stochastic processes (dispersal limitation, drift, and homogenizing dispersal) that govern the assembly of the phyllosphere bacterial communities. Drift (25–54%), dispersal limitation (10–52%), and homogeneous selection (15–32%) were the dominant assembly processes across months and canopy positions (Figure 4A) (File S5). A total of 126 bins representing ASVs that are phylogenetically related and putatively share similar ecological functions were detected in this study. The 15 top bins accounting for 62% of the total relative abundance were dominated by ASVs affiliated with *Sphingomonas*, unclassified *Beijerinckiaceae, Pseudomonas*, *Methylobacterium–Methylorubrum, Rosenbergiella*, *Curtobacterium*, *Pantoea*, *Massilia*, and *Erwinia*) (File S6).

**Figure 4.**
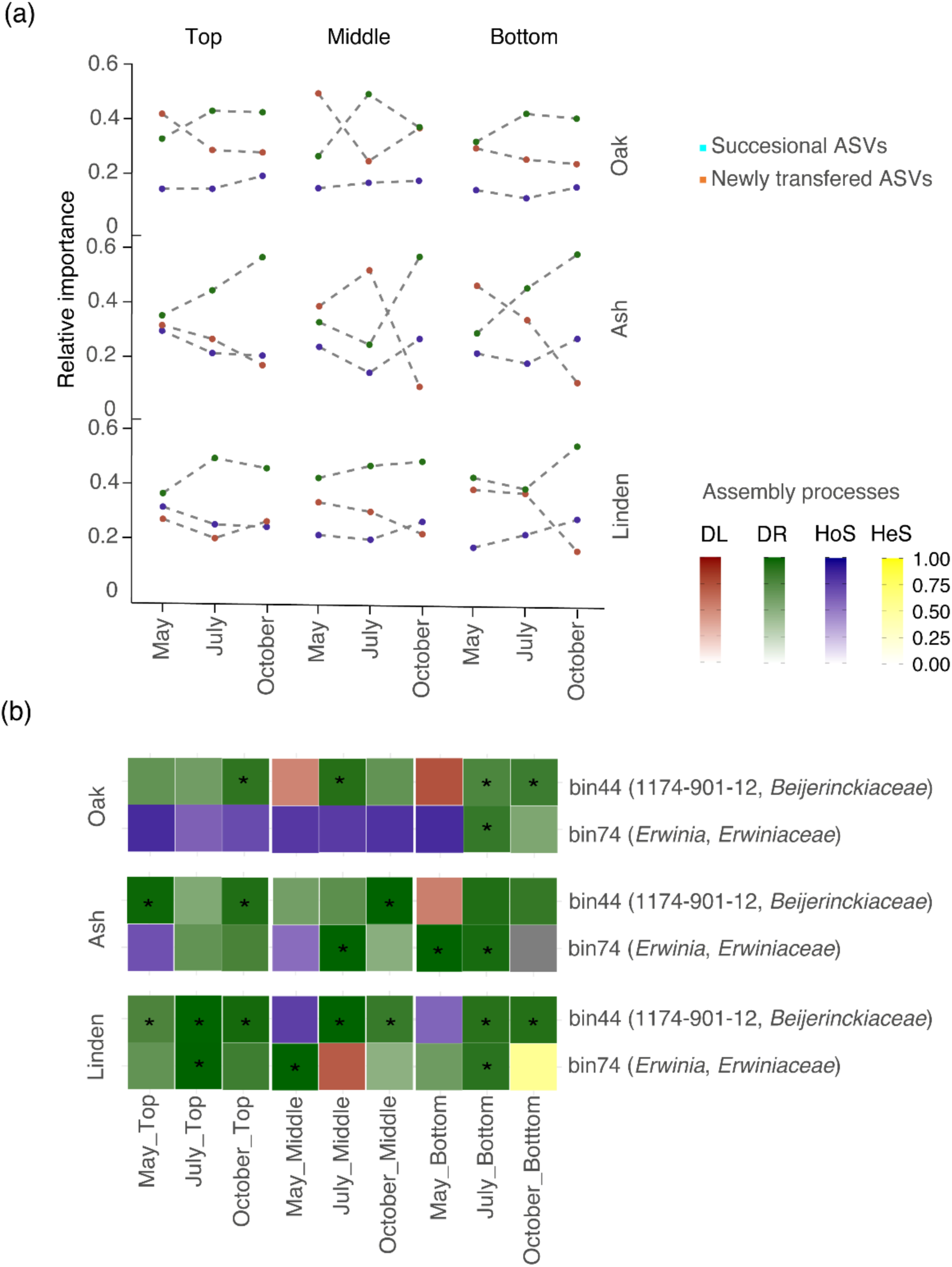
Relative importance of assembly process and key actual sequence variants across canopy positions, phenological stages and tree species. The relative importance (0 to 1) of assembly processes across canopy positions, phenological stages and tree species for (a) overall and (b) key bins. * denotes statistical significance (*P*<0.05) of the process contributed by the respective bins as per iCAMP analysis.

At an early phenological stage, community assembly was dominated by dispersal limitation (32–47%), drift (29–38%), and homogenous selection (17–30%), which shifted towards drift (25–58%) during the mid and late phenological stages for oak and ash (Figure 4A). For linden, the drift–based community assembly dominated across canopy positions and phenological stages (37–54%) (Figure 4A). The contribution of successional and newly introduced ASVs towards the three dominant processes (homogenous selection, drift and dispersal limitation) showed that the successional ASVs were the major driver of drift, while the newly introduced ASVs were linked to dispersal limitation (Figure 4A). As the communities transitioned from early to mid and late phenological stages, the relative importance of dispersal limitation decreased which was linked to decreasing abundances of *Erwiniaceae* (bin 74), while the relative importance of drift mediated by *Beinjerinckiaceae* (bin 44) increased (Figure 4B and File S5).

### Functional potential of tree phyllosphere bacteria across the phenological stages

Functional potential analysis of phyllosphere bacteria identified 65 level 3 KEGG pathways associated with the category “metabolism”, of which the majority remained unchanged between mid and late phenological stages (Kruskal–Wallis and Dunn’s pairwise test, P > 0.05) across all tree species (64 in oak, 58 in ash, and 60 in linden)(Figure 5 and File S7). For all three tree species, KEGG pathways belonging to type II polyketide synthase (PKS) and steroid and sphingolipid biosynthesis pathways belonging to lipid metabolism were enriched during mid and late phenological stages (Kruskal–Wallis and Dunn’s pairwise test, *P*<0.05). Additionally, we observed an increase in flavonoid, butirosin and neomycin biosynthesis pathway abundances (ash), flavonoid biosynthesis pathways (linden), and non-ribosomal peptide pathways (oak) during the mid and late phenological stages (Kruskal–Wallis and Dunn’s pairwise test, *P*<0.05).

**Figure 5.**
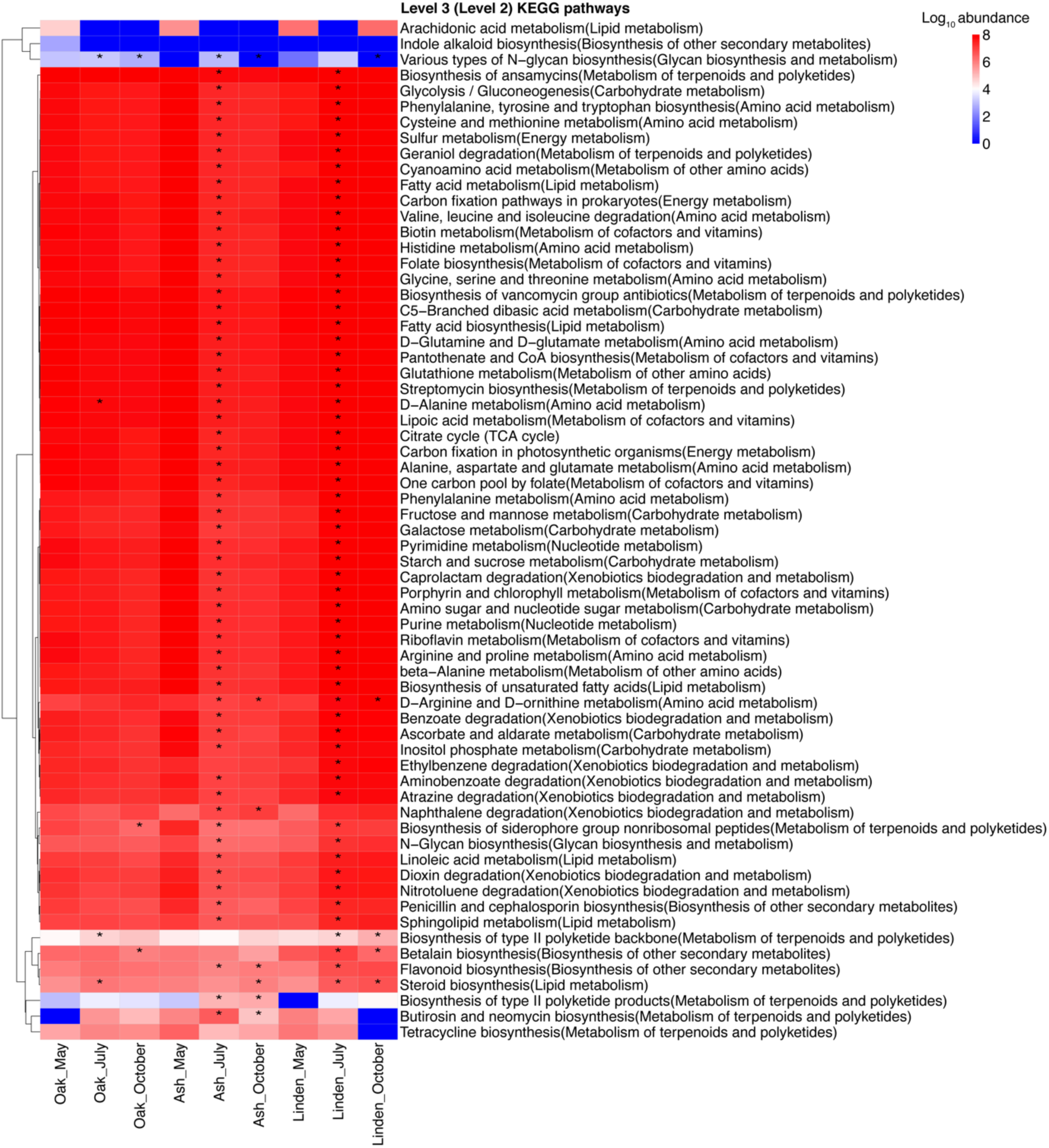
Heatmap showing KEGG Level 2 and Level 3 pathway abundances at phenological stage and tree species level. The Euclidean based hierarchical clustering was applied for rows (KEGG pathway abundances). The colour intensity of the palette indicates the log transformed pathway abundances calculated from estimated cell abundances (g^-1^ dry leaf) from low (blue) to high (red). * in July and October indicates statistical significance (Kruskal–Wallis and Dunn’s pairwise test, *P*<0.05) when compared with their respective preceding phenological stage.

### Bacterial communities in throughfall

Similar to the phyllosphere communities, phenological stages explained 19.98% of the variation in bacterial communities transported by throughfall, followed by the influencing factors tree species (4.38%), tree individual (3.89%) and canopy position (2.22%) (Figure 6A, Table S2). Throughfall bacterial communities collected in March before leaf sprouting were distinct from the other phenological stages. In contrast to the phyllosphere communities, tree individuals were the third best factor explaining variation in throughfall communities (Table S2).

**Figure 6.**
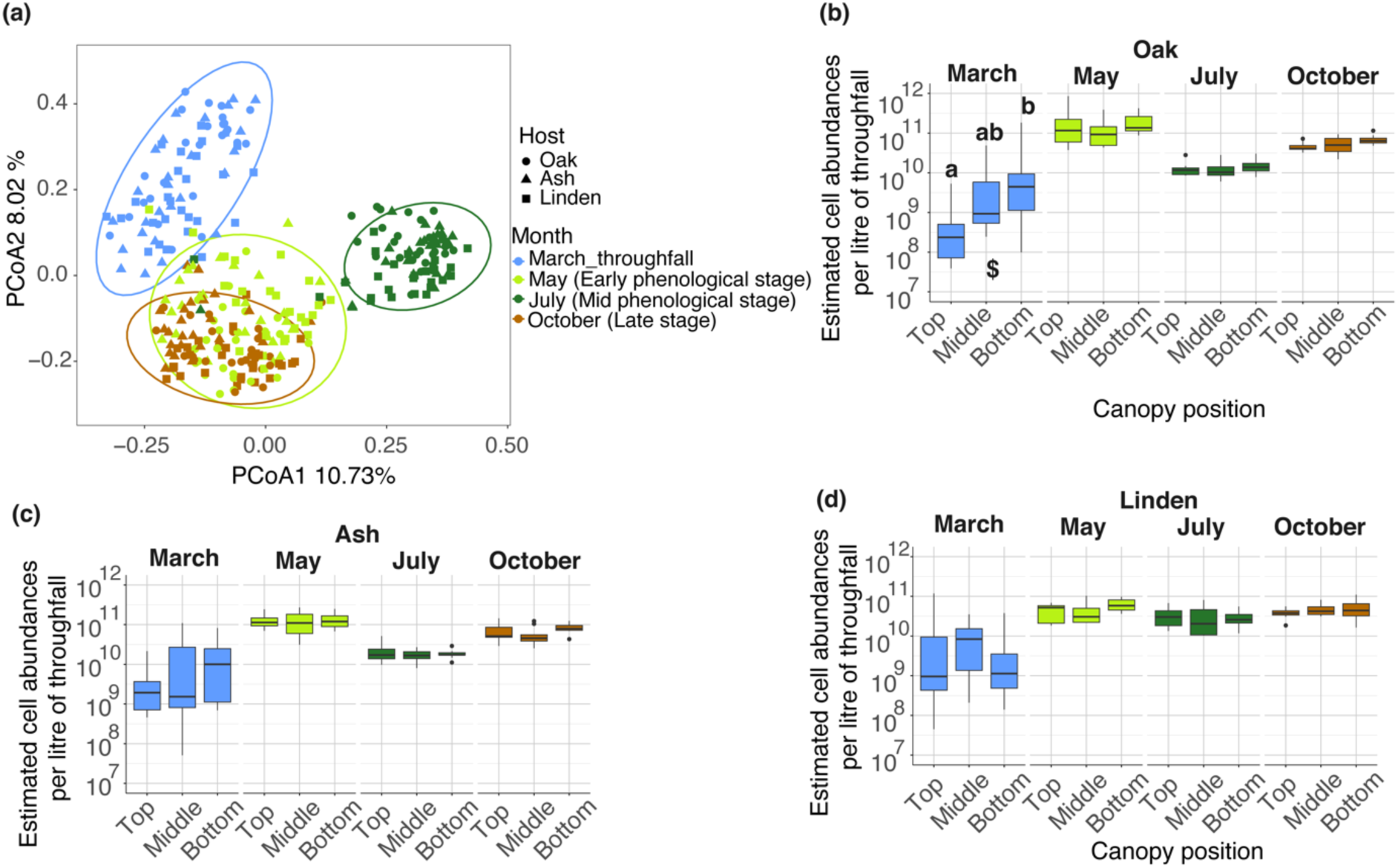
Principal co-ordinate analysis of throughfall bacterial communities associated with oak, ash and linden for the three phenological stages (early, mid and late phenological stages) and additionally for the leafless stage in March. (a) Sample-based ordination plot generated from amplicon sequence variant (ASVs) (operon normalised) compositions of throughfall bacteria. Ellipses represent 95% confidence intervals for the tested factor variable (i.e. Phenological stages), (b-d) Estimated cell abundances of throughfall bacteria across tree species and canopy positions. Alphabets above the boxplots denote statistical difference between canopy positions. ^$^ below the boxplots indicates depicts statistical significance (*P*<0.05) based on Kruskal’s wallis and Dunn’s pairwise test (*P*<0.05).

The Shannon’s diversity index H’ was highest for throughfall collected in May and July across all tree species (Figure S6). At canopy position level, throughfall Shannon’s diversity index H’ was highest in bottom positions during the early and mid-phenological stage for ash and linden, and during early and late phenological stages for oak (Figure S6). There was no significant difference of Shannon’s diversity index H’ for throughfall-affiliated bacterial communities between oak, ash and linden. Estimated bacterial cell abundances in throughfall ranged from 1.1×10^9^ to 2.2×10^11^ per liter across all tree species and time points (File S2) with lowest abundances in throughfall collected in March, and higher abundances during the mid and late phenological stages (Kruskal–Wallis and Dunn’s pairwise test, *P<*0.05) (Figure 6B–6D).

In order to estimate the total flux of bacteria being transported by rainwater through the canopy system, we calculated bacterial fluxes per m^2^ and sampling event. Throughfall bacterial flux remained unchanged across canopy positions for oak, whereas for ash and linden, the mean flux was highest at the top (1.9×10^12^ or 2.5×10^12^ bacterial cells m^-2^ sampling event^-1^) compared to the bottom (8×10^11^ or 1.07×10^12^ bacterial cells m^-2^ sampling event^-1^) canopy position during the mid–phenological stage (Kruskal–Wallis and Dunn’s pairwise test, *P<*0.05) (Figure S7 and Table S6). For linden, a similar trend was also observed during the late phenological stage.

The bacterial communities transported by throughfall consisted of families that were also commonly detected in the phyllosphere, including *Pseudomonadaceae*, *Oxalobacteraceae*, *Microbacteraceae*, *Beijerinckiaceae*, *Erwiniaceae*, *Rhizobiaceae* and *Spirosmaceae* (Figure S8)*. Oxalobacteraceae* and *Pseudomonadaceae* clearly dominated the throughfall communities in March before foliage emerged. While ASVs affiliated with *Sphingomonas*, *Pseudomonas*, and *Massilia* exhibited consistently high abundances in throughfall across all tree species and time points, relative abundances of ASVs affiliated with *Methylobacterium-Methylorubrum*, *Kineococcus*, *Acidiphilum*, *Curtobacterium*, *Pantoea*, and *Erwinia* increased only after the sprouting of leaves in May (Figure 7A). Estimated throughfall-mediated vertical transfer pointed to a lower exchange of ASVs between throughfall and phyllosphere surfaces in the mid (5–21%) compared to the early (11–23%) and late phenological stages (13–26%) (Figure 7B) (Kruskal–Wallis and Dunn’s pairwise test, *P<*0.05).

**Figure 7.**
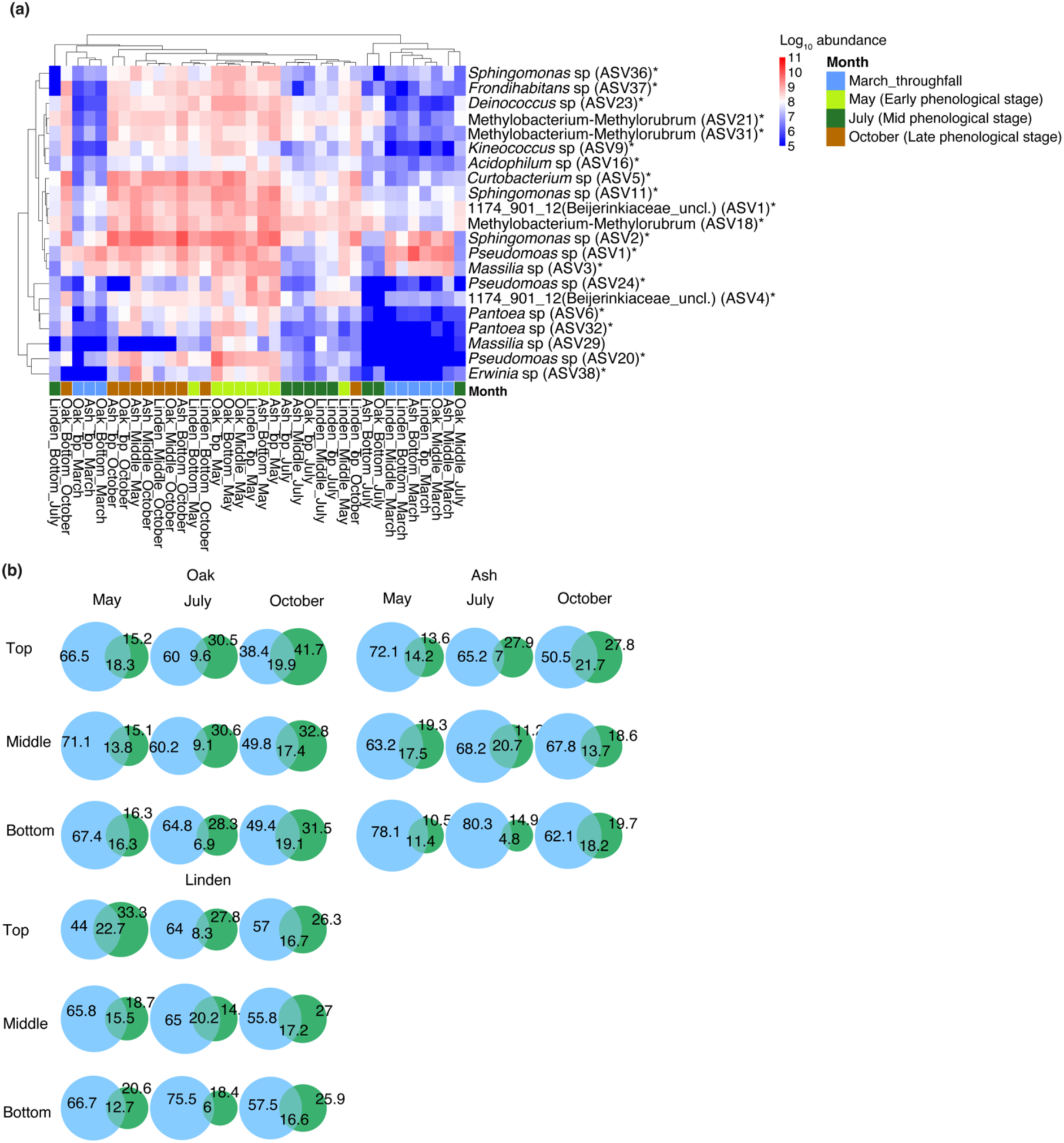
Top throughfall amplicon sequence variants and shared amplicon sequence variants (throughfall–phyllosphere) across tree species, phenological stages and canopy positions. (a) Heatmap showing top throughfall amplicon sequence variants (ASVs). The colour intensity of the palette indicates the log transformed estimated cell abundances from low (blue) to high (red). (b) Venn diagram showing the shared and unique amplicon sequence variant (ASV) percentages between throughfall and phyllosphere samples for early, mid and late phenological stages. March-throughfall indicates throughfall samples before the start of bud bursting. * denotes differentially abundant (*P*<0.05) throughfall ASVs based on LEfSe analysis.

Bacterial taxa prone for either leaf attachment in the phyllosphere or wash-off during successional transfer (early, mid and late phenological stages) and vertical transfer (crane top, top, mid and bottom canopy positions) were identified based on LDA scores (LEfSe analysis) derived from the relative abundance of shared ASVs in throughfall versus phyllosphere (*P<*0.05 and FDR <0.05) (Figure 8A and File S4). A consistent higher prevalence in the phyllosphere was observed for ASVs affiliated with *Kineococcus* sp, *Pseudomonas* sp, *Massilia* sp, *Hymenobacter* sp, *Pantoea* sp., *Sphingomonas*, *Methylobacterium-Methylorubrum*, *Methylocella*, and other *Beijerinckiaceae*, *Microbacteriacae*, and *Acetobacteraceae* (Figure 8B). In contrast, a higher tendency to get mobilized by throughfall was detected for ASVs affiliated with *Allo*-*Neo-Para-rhizobium* and *Deinococcus* across all tree species, or for *Janthinobacterium* and *Novosphingobium* only in association with oak or linden, respectively (Figure 8B).

**Figure 8.**
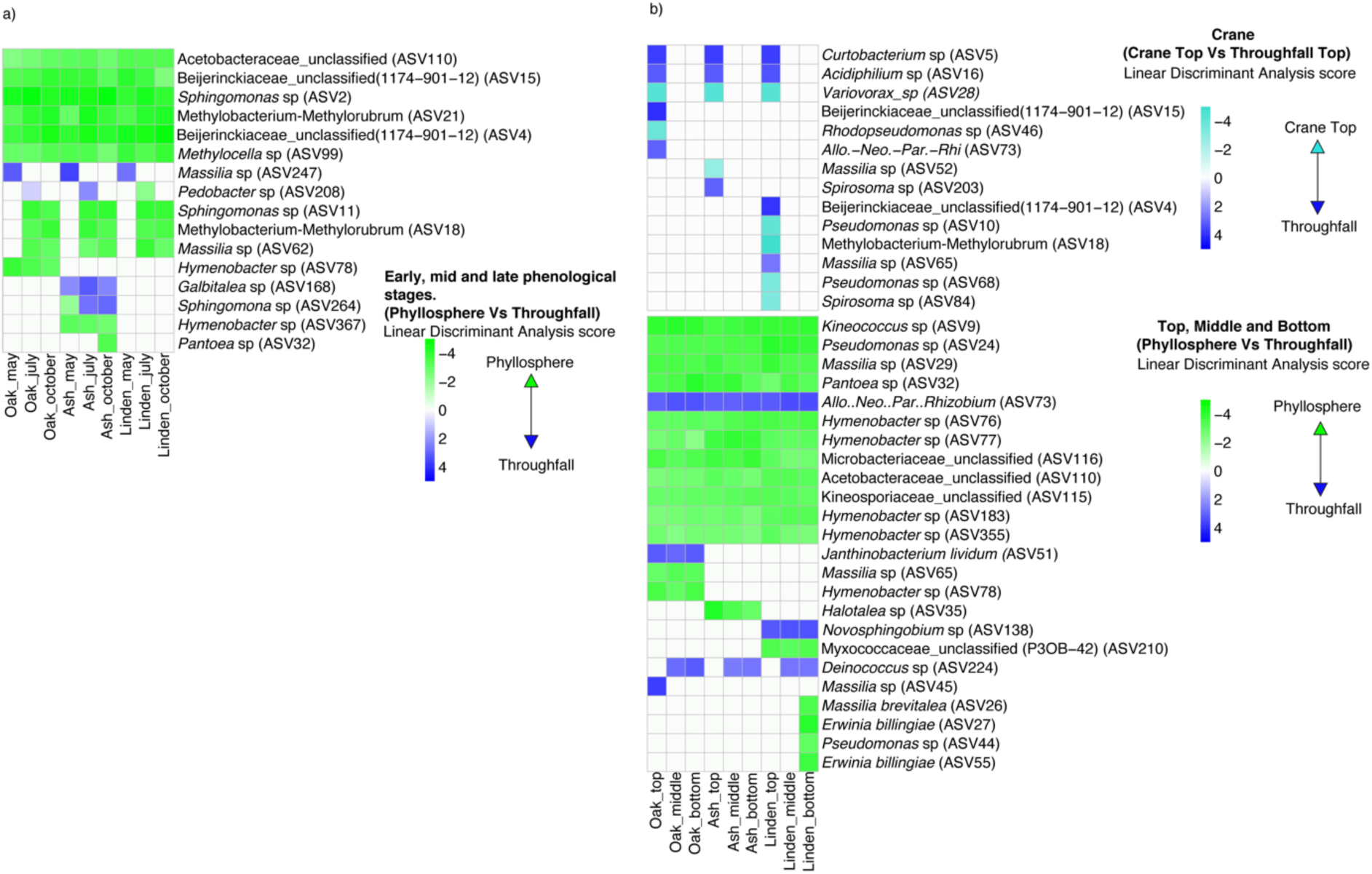
Successional and vertical trends of phyllosphere attachment/throughfall wash–off patterns of amplicon sequence variants. The relative abundances of common ASVs between groups (crane top vs throughfall or phyllosphere vs throughfall) were analyzed at a) successional (early, mid and late phenological stages) and b) vertical levels (crane top, top, middle and bottom canopy positions). LDA scores were calculated using LEfSe (*P*<0.05 and FDR<0.05). For crane top vs throughfall top comparisons, turquoise blue color indicates crane top enrichment, blue indicates throughfall wash-off at top position. For phyllosphere vs throughfall comparisons, green color indicates preferential attachment, blue indicates preferential throughfall-mediated wash-off across all canopy positions.

## Discussion

Our findings clearly demonstrated that phenological stage is a key driver of phyllosphere microbiome variability in a temperate hardwood forest, vastly exceeding the effects of tree species identity. In contrast to our expectations and earlier observations (Herrmann et al. 2021), canopy position explained only a minor part of phyllosphere microbiome variability. Yet, we present strong evidence that canopy-position dependent vertical trends of phyllosphere bacterial abundance are tightly coupled to throughfall-mediated within-canopy transport. While phenology was previously recognized to play a key role in phyllosphere microbiome variability in evergreen (*Pinus* and *Sabina*) (Bao et al., 2020) and perennial (*Citrus* and *Acacia*) plants (Al Ashhab et al., 2021; Ginnan et al., 2022), our observation is in contrast to earlier studies of temperate forest ecosystems that primarily identified host taxonomic and genetic identity as the main drivers of phyllosphere bacterial communities (Laforest-Lapointe et al., 2017; Wang et al., 2023).

Early phenological stage communities after leaf sprouting were distinct, while phyllosphere bacterial communities became more similar and functionally redundant during mid–late phenological stages. The observed phenological pattern may have been caused by combined effects of tree species specific factors and external factors including leaf-visiting insects for pollination or herbivory (Ekholm et al., 2020), weather conditions, and inputs of pollen and dust. From the tree host perspective, phenological stages are characterized by changes that synchronize with bud burst and leaf out at the early phenological stage to heightened metabolic activities during mid–late phenological stages such as photosynthesis (Wong et al., 2020), changes in leaf chemistry (Oktavia and Jin, 2020) and trace gas production (Vermeuel et al., 2023).

In line with these phenology-driven changes, phyllosphere communities in July and October shared a larger fraction of successional ASVs compared to the fraction shared with the early phenological stage in May, suggesting a transition from bacterial communities which are mostly characterized by random colonization to communities that are more closely associated with tree host properties, as the vegetation period progressed. In fact, the fraction of phyllosphere bacterial community variability explained by tree species identity increased from 19.48% in May to 28.37% in October, indicating an increasing relevance of tree-species specific drivers with ongoing leaf maturation and senescence. Young leaves are known to have a higher nitrogen content (Feeny, 1970; Hoffland et al., 2000; Mediavilla and Escudero, 2003). Consequently, early bacterial colonizers likely occupied leaf buds and quickly depleted the available resources including nitrogen compounds through competitive exclusion (Posfai et al., 2017), thereby repelling the colonisation by other bacteria during the early phenological stages. Increased availability of leaf-associated resources such as volatile organic compounds, sugars, or amino acids during mid and late phenological stages (Tukey, 1970, Van Stan et al., 2012, Chen and Poland, 2009) likely increased phyllosphere niche diversity and enabled the growth, establishment, co-existence, and successional transfer of more taxa (Posfai et al., 2017), resulting in the observed diverse phyllosphere bacterial communities of the mid and late phenological stages. The quantitative relevance of newly introduced bacteria was typically lower during the late phenological stage as the phyllosphere habitat became increasingly colonized and occupied.

The described phenology-driven changes in host plant physiology as well as seasonal immigration patterns of atmospheric, pollen- or insect-associated microorganisms are likely to have a strong effect on the competition between introduced and successional bacterial leaf colonizers. In turn, the balance between introduction, competition and persistence of microbial taxa may be crucial for maintaining necessary ecosystem functions at the respective phenological stages. While moving from early to mid and late phenological stages, we noted a shift of bacterial communities from being dominated by putative plant pathogens (e.g. *Erwiniaceae*) to being dominated by bacterial taxa associated with plant growth promotion and biocontrol (e.g. *Beijerinckiaceae*) (Tamas et al., 2014). Low plant defense capabilities (Sancho-Knapik et al., 2023) are characteristic of plant physiological conditions at early phenological stages, which possibly favoured the colonization by bacterial pathogens such as *Erwiniaceae*. The observed later decrease in the abundance of putative plant pathogens (*Erwinia* sp., *Pseudomonas* sp., *Pantoea* sp., and *Serratia* sp.) during mid and late phenological stages likely occurred due to competitive exclusion by early colonizers (Maignien et al., 2014; Schlechter et al., 2023). The higher abundance of putative steroid and secondary metabolite related pathways (Type II PKS, flavonoid and non-ribosomal peptide) indicates the importance of signalling molecules towards plant growth promotion (Ai et al., 2023) and antagonistic mechanisms in pathogen control (Bertić et al., 2023), respectively, during mid-late phenological stages.

Despite the increasing influence of tree species on phyllosphere communities as the vegetation period progressed, bacterial community assembly in the phyllosphere was mainly driven by stochastic processes, especially dispersal limitation and drift. In models of stochastic assembly, random factors or events, including lack of movement, colonization and extinction shape community composition over time (Zhou and Ning, 2017). Dispersal limitation dominated during the early phenological stage, likely due to the spatial separation of small leaf sprouts and the random nature of the colonization events. The early colonizers, especially pathogens (e.g. *Erwinia* sp), likely depleted the available space and resources during leaf budding through priority effects (Maignien et al., 2014). Similarly, previous work identified priority effects as a driving factor for heterogeneous phyllosphere bacterial communities, along with a dominance of dispersal limitation at an early phenological stage (Maignien et al., 2014). During mid and late phenological stages, new bacteria entered and shared the resources with successional bacteria (e.g. *Beijerinkiacea*e) through niche partitioning, which led to a shift of the dominant assembly process towards drift. Despite continued taxonomic turnover, the higher abundance of bacteria carrying out methylotrophic and metabolically diverse functions in the communities of the mid and late phenological stages resulted in functional redundancy (Ren et al., 2020), which has been reported both as a consequence as well as a potential driving factor for drift assembly (Li et al., 2021; Ren et al., 2020). Functional redundancy was further reflected in the stable abundance patterns across predicted homeostasis pathways, e. g., carbohydrate and energy metabolism.

Assessment of rainwater-mediated bacterial transport into and within the canopy system allowed us to identify potential airborne import of microbes and taxon-specific mobilization patterns. In general, throughfall-mediated transport and mobilization resulted in a substantial flux of microbial cells through the tree crowns of up to 10^12^ cells per m^2^ per sampling event, which is in the same range as throughfall bacterial loads measured below the canopy in a previous study (Krüger et al., 2025). These findings support our hypothesis that rainwater plays a key role in the distribution of bacteria within the canopy, acting as an important facilitator of bacterial colonization but also detachment. Contact of throughfall with foliage resulted in an increase of transported bacteria by up to two orders of magnitude when comparing leafless (March) and foliated stage (May), or rainwater collected above the tree canopy to throughfall collected at the canopy top.

For both oak and linden, Shannon diversity and abundances of the phyllosphere bacteria were highest at the bottom position of the canopy, suggesting that throughfall-mediated wash-off and transport from above parts resulted in an increased bacterial import to the lowest position of the tree crown. Throughfall-mediated mobilization was the most intense at the canopy top across all three tree species, coinciding with lowest phyllosphere bacterial diversity and abundance at this position. Consequently, reduced diversity and abundance at the canopy top might not only be a result of generally harsher conditions at this exposed position (Duan et al., 2024), but phyllosphere taxa might also be more prone to detachment under these conditions. This likely results in continuous transfer of microbial biomass to the mid and bottom canopy where reduced physical impact of rainfall (Stone and Jackson 2019), more favorable microclimatic conditions, and higher nutrient availability might allow a denser colonization with more efficient protection against detachment. Despite the strong effect on vertical phyllosphere abundance and diversity patterns, transport-related mechanisms did not result in consistent canopy position dependent community patterns across all tree species and phenological stages, as canopy position only explained a maximum of 6% of total community variation. The fact that canopy position contributed 15% of community variation in September 2017 at the same study site (Herrmann et al. 2021) points to an important role of the specific weather conditions of a given year for vertical patterns of the phyllosphere microbiome.

The vertical distribution of individual bacterial taxa appeared to be strongly influenced by potential airborne import and attachment and detachment patterns during exchange between throughfall and phyllosphere. Taxa typically associated with cloud water or atmospheric inputs such as *Massilia* (Li et al., 2020; Woo and Yamamoto, 2020; Zhang et al., 2022) or *Pseudomonas* (Woo and Yamamoto, 2020) were abundant in rainwater above the canopy and in throughfall but exhibited a strong tendency to preferentially occur in the phyllosphere, pointing to strong adhesion mechanisms. In fact, 10–40% of taxa typically occurring in the phyllosphere are known to form biofilms for attachment (Morris et al., 1998), including the genera *Hymenobacter* sp., *Massilia* sp., and *Pseudomonas* sp. which dominated the phyllosphere communities in this study. A stronger tendency for an attached lifestyle of *Massilia* sp. in association with oak, *Halotalea* sp. in association with ash, and unclassified *Myxococcaceae* in association with linden, further pointed to a central role of the host tree in selectively favoring attachment versus wash-off of individual taxa.

Similar to the phyllosphere communities, we observed a strong seasonal imprint on throughfall bacterial community composition. Throughfall collected in March during the leafless stage of the canopy harboured the most distinct bacterial communities, suggesting that microbial exchange between throughfall and phyllosphere after foliage emerged had a strong converging effect on the transported communities. Notably, tree individual played a much more important role in explaining throughfall than phyllosphere bacterial community variability, likely resulting from the effect of the unique branching architecture of individual trees on water distribution patterns and thus also microbial transport within the individual tree crowns (MacFarlane and Kane, 2017; Su et al., 2019).

### Conclusions

Our findings revealed a strong impact of phenological stages on bacterial communities and their assembly in the phyllosphere of temperate forests, overriding tree species-specific or canopy position dependent effects. Yet, an increasing influence of tree species identity towards mid and late phenological stages, along with increased functional redundancy, suggested a transition from phyllosphere communities which are mostly controlled by random colonization events and dispersal limitation in spring towards communities which are increasingly subject to tree-species specific controls. Our unique experimental setup further identified throughfall as a key mechanism importing and transporting bacteria through the tree crowns with taxon-specific mobilization patterns, which likely underlies the observed maxima of diversity and phyllosphere bacterial abundance at the bottom of the canopy. Our work underscores the need to integrate phenological aspects as well as the whole vertical extension of tree canopies to understand phyllosphere microbiome dynamics in temperate forests.

## Acknowledgements

We thank Stefan Riedel, Jens Wurlitzer and Falko Gutmann for help with field work and sample processing. René Maskos and Stefan Riedel are acknowledged for excellent technical support with amplicon library preparation and sequencing. We thank Martin Stephan, head of the mechanics workshop at Otto-Schott-Institute, FSU Jena, for manufacturing the customized throughfall samplers. This study was financially supported by the German Center for Integrative Biodiversity Research (iDiv) Halle-Jena-Leipzig, funded by the Deutsche Forschungsgemeinschaft (DFG, German Research Foundation) – Project Number FZT 118. Additional financial support was provided by the Collaborative Research Center 1076 AquaDiva (Deutsche Forschungsgemeinschaft, project number 218627073). The infrastructure for MiSeq Illumina sequencing was financially supported by the Thüringer Ministerium für Wirtschaft, Wissenschaft und Digitale Gesellschaft (TMWWDG; project 2016 FGI 0024 “BIODIV”).

## Author contributions

DSLC performed all sequence data and ecological analysis and wrote the manuscript with contributions from all other co-authors. MH, BM and CW designed the study and acquired funding. BM designed the approach of within-canopy throughfall sampling. MMS, PZ and MH performed all the molecular work. RR, RE and TK provided access to the canopy crane platform, planned the field work, operated the crane, and provided field and climate data. All authors contributed to writing of the manuscript.

## Data availability statement

The amplicon sequencing datasets generated in this study have been deposited in the European Nucleotide Archive (ENA) under the BioProject IDs PRJEB109949 and PRJEB109950. Results of quantitative PCR, PICRUSt analysis and community assembly analysis are provided as Supplementary Material. R Codes used in this work are provided at Github (link).

## References

Ai, W., Liu, H., Wang, Yutao, Wang, Yu, Wei, J., Zhang, X., Lu, X., 2023. Identification of Functional Brassinosteroid Receptor Genes in Oaks and Functional Analysis of QmBRI1. Int. J. Mol. Sci. 24, 16405. 10.3390/ijms242216405

Al Ashhab, A., Meshner, S., Alexander-Shani, R., Dimerets, H., Brandwein, M., Bar-Lavan, Y., Winters, G., 2021. Temporal and Spatial Changes in Phyllosphere Microbiome of Acacia Trees Growing in Arid Environments. Front. Microbiol. 12. 10.3389/fmicb.2021.656269

Bao, L., Gu, L., Sun, B., Cai, W., Zhang, S., Zhuang, G., Bai, Z., Zhuang, X., 2020. Seasonal variation of epiphytic bacteria in the phyllosphere of Gingko biloba, Pinus bungeana and Sabina chinensis. FEMS Microbiol. Ecol. 96. 10.1093/femsec/fiaa017

Bechtold, E.K., Wanek, W., Nuesslein, B., DaCosta, M., Nüsslein, K., 2024. Successional changes in bacterial phyllosphere communities are plant-host species dependent. Appl. Environ. Microbiol. 10.1128/aem.01750-23

Bertić, M., Orgel, F., Gschwendtner, S., Schloter, M., Moritz, F., Schmitt-Kopplin, P., Zimmer, I., Fladung, M., Schnitzler, J., Schroeder, H., Ghirardo, A., 2023. European oak metabolites shape digestion and fitness of the herbivore Tortrix viridana. Funct. Ecol. 37, 1476–1491. 10.1111/1365-2435.14299

Bittar, T.B., Pound, P., Whitetree, A., Moore, L.D., Van Stan, J.T., 2018. Estimation of Throughfall and Stemflow Bacterial Flux in a Subtropical Oak-Cedar Forest. Geophys. Res. Lett. 45, 1410–1418. 10.1002/2017GL075827

Callahan, B.J., McMurdie, P.J., Rosen, M.J., Han, A.W., Johnson, A.J.A., Holmes, S.P., 2016. DADA2: High-resolution sample inference from Illumina amplicon data. Nat. Methods 13, 581–583. 10.1038/nmeth.3869

Carrell, A.A., Frank, A.C., 2014. Pinus flexilis and Piceae engelmannii share a simple and consistent needle endophyte microbiota with a potential role in nitrogen fixation. Front. Microbiol. 5. 10.3389/fmicb.2014.00333

Chase, J.M., Myers, J.A., 2011. Disentangling the importance of ecological niches from stochastic processes across scales. Philos. Trans. R. Soc. B Biol. Sci. 366, 2351–2363. 10.1098/rstb.2011.0063

Chelius, M.K., Triplett, E.W., 2001. The diversity of archaea and bacteria in association with the roots of Zea mays L. Microb. Ecol. 41, 252–263. 10.1007/s002480000087

Chen, D., Yang, J., Wang, S., Lan, S., Wang, Y., Liu, Z.-J., Qian, X., 2025. Comparative analysis of community composition and network structure between phyllosphere endophytic and epiphytic fungal communities of Mussaenda pubescens. Microbiol. Spectr. 13. 10.1128/spectrum.01019-24

Chen, Y., Poland, T.M., 2009. Interactive Influence of Leaf Age, Light Intensity, and Girdling on Green Ash Foliar Chemistry and Emerald Ash Borer Development. J. Chem. Ecol. 35, 806–815. 10.1007/s10886-009-9661-1

De Mandal, S., Jeon, J., 2023. Phyllosphere Microbiome in Plant Health and Disease. Plants 12, 3481. 10.3390/plants12193481

Deng, N., Liu, C., Tian, Y., Song, Q., Niu, Y., Ma, F., 2024. Assembly processes of rhizosphere and phyllosphere bacterial communities in constructed wetlands created via transformation of rice paddies. Front. Microbiol. 15. 10.3389/fmicb.2024.1337435

Douglas, G.M., Maffei, V.J., Zaneveld, J.R., Yurgel, S.N., Brown, J.R., Taylor, C.M., Huttenhower, C., Langille, M.G.I., 2020. PICRUSt2 for prediction of metagenome functions. Nat. Biotechnol. 38, 685–688. 10.1038/s41587-020-0548-6

Duan, Y., Siegenthaler, A., Skidmore, A.K., Chariton, A.A., Laros, I., Rousseau, M., De Groot, G.A., 2024. Forest top canopy bacterial communities are influenced by elevation and host tree traits. Environ. Microbiome 19, 21. 10.1186/s40793-024-00565-6

Ekholm, A., Tack, A.J.M., Pulkkinen, P., Roslin, T., 2020. Host plant phenology, insect outbreaks and herbivore communities – The importance of timing. J. Anim. Ecol. 89, 829–841. 10.1111/1365-2656.13151

Feeny, P., 1970. Seasonal Changes in Oak Leaf Tannins and Nutrients as a Cause of Spring Feeding by Winter Moth Caterpillars. Ecology 51, 565–581. 10.2307/1934037

Ginnan, N.A., De Anda, N.I., Campos Freitas Vieira, F., Rolshausen, P.E., Roper, M.C., 2022. Microbial Turnover and Dispersal Events Occur in Synchrony with Plant Phenology in the Perennial Evergreen Tree Crop Citrus sinensis. MBio 13. 10.1128/mbio.00343-22

Hassan, A.., Frank, J.., 2003. Influence of surfactant hydrophobicity on the detachment of Escherichia coli O157:H7 from lettuce. Int. J. Food Microbiol. 87, 145–152. 10.1016/S0168-1605(03)00062-X

Herrmann, M., Geesink, P., Richter, R., Küsel, K., 2021. Canopy Position Has a Stronger Effect than Tree Species Identity on Phyllosphere Bacterial Diversity in a Floodplain Hardwood Forest. Microb. Ecol. 81, 157–168. 10.1007/s00248-020-01565-y

Hoffland, E., Jeger, M.J., Van Beusichem, M.L., 2000. Effect of nitrogen supply rate on disease resistance in tomato depends on the pathogen. Plant Soil 218, 239–247.

Iguchi, H., Yurimoto, H., Sakai, Y., 2015. Interactions of Methylotrophs with Plants and Other Heterotrophic Bacteria. Microorganisms 3, 137–151. 10.3390/microorganisms3020137

Jansen, E., 1999. Das Naturschutzgebiet Burgaue. Staatliches Umweltfachamt, Leipzig.

Jiao, S., Lu, Y., 2020. Soil pH and temperature regulate assembly processes of abundant and rare bacterial communities in agricultural ecosystems. Environ. Microbiol. 22, 1052–1065. 10.1111/1462-2920.14815

Klindworth, A., Pruesse, E., Schweer, T., Peplies, J., Quast, C., Horn, M., Glöckner, F.O., 2013. Evaluation of general 16S ribosomal RNA gene PCR primers for classical and next-generation sequencing-based diversity studies. Nucleic Acids Res. 41, e1. 10.1093/nar/gks808

Krüger, M., Potthast, K., Michalzik, B., Tischer, A., Herrmann, M., 2025. Rainwater-driven transport of matter and microbes from phyllosphere to soil in a temperate beech forest. Sci. Total Environ. 1006, 180910. 10.1016/j.scitotenv.2025.180910

Krüger, M., Potthast, K., Michalzik, B., Tischer, A., Küsel, K., Deckner, F.F.K., Herrmann, M., 2021. Drought and rewetting events enhance nitrate leaching and seepage-mediated translocation of microbes from beech forest soils. Soil Biol. Biochem. 154, 108153. 10.1016/j.soilbio.2021.108153

Laforest-Lapointe, I., Messier, C., Kembel, S.W., 2017. Tree Leaf Bacterial Community Structure and Diversity Differ along a Gradient of Urban Intensity. mSystems 2. 10.1128/mSystems.00087-17

Laforest-Lapointe, I., Messier, C., Kembel, S.W., 2016. Host species identity, site and time drive temperate tree phyllosphere bacterial community structure. Microbiome 4, 27. 10.1186/s40168-016-0174-1

Laforest-Lapointe, I., Whitaker, B.K., 2019. Decrypting the phyllosphere microbiota: progress and challenges. Am. J. Bot. 106, 171–173. 10.1002/ajb2.1229

Levia, D.F., Keim, R.F., Carlyle-Moses, D.E., Frost, E.E., 2011. Throughfall and Stemflow in Wooded Ecosystems. pp. 425–443. 10.1007/978-94-007-1363-5_21

Li, Q., Long, Z., Wang, H., Zhang, G., 2021. Functions of constructed wetland animals in water environment protection – A critical review. Sci. Total Environ. 760, 144038. 10.1016/j.scitotenv.2020.144038

Li, T., Gao, J., 2023. Attribution of dispersal limitation can better explain the assembly patterns of plant microbiota. Front. Plant Sci. 14. 10.3389/fpls.2023.1168760

Li, X., Chen, H., Yao, M., 2020. Microbial emission levels and diversities from different land use types. Environ. Int. 143, 105988. 10.1016/j.envint.2020.105988

Lusk, M.G., 2024. Throughfall as an understudied biogeochemical subsidy of nutrients and carbon in the urban water cycle: perspective and a research agenda. Discov. Water 4, 124. 10.1007/s43832-024-00181-y

Lyautey, E., Jackson, C.R., Cayrou, J., Rols, J.-L., Garabétian, F., 2005. Bacterial Community Succession in Natural River Biofilm Assemblages. Microb. Ecol. 50, 589–601. 10.1007/s00248-005-5032-9

MacFarlane, D.W., Kane, B., 2017. Neighbour effects on tree architecture: functional trade-offs balancing crown competitiveness with wind resistance. Funct. Ecol. 31, 1624–1636. 10.1111/1365-2435.12865

Maignien, L., DeForce, E.A., Chafee, M.E., Eren, A.M., Simmons, S.L., 2014. Ecological Succession and Stochastic Variation in the Assembly of Arabidopsis thaliana Phyllosphere Communities. MBio 5. 10.1128/mBio.00682-13

Mediavilla, S., Escudero, A., 2003. Relative growth rate of leaf biomass and leaf nitrogen content in several mediterranean woody species. Plant Ecol. 168, 321– 332. 10.1023/A:1024496717918

Morris, C.E., Kinkel, L., 2002. Fifty years of phyllosphere microbiology: Significant contributions to research in related fields, in: Phyllosphere Microbiology. pp. 365–375.

Morris, C.E., Monier, J.-M., Jacques, M.-A., 1998. A Technique To Quantify the Population Size and Composition of the Biofilm Component in Communities of Bacteria in the Phyllosphere. Appl. Environ. Microbiol. 64, 4789–4795. 10.1128/AEM.64.12.4789-4795.1998

Ning, D., Yuan, M., Wu, L., Zhang, Y., Guo, X., Zhou, X., Yang, Y., Arkin, A.P., Firestone, M.K., Zhou, J., 2020. A quantitative framework reveals ecological drivers of grassland microbial community assembly in response to warming. Nat. Commun. 11, 4717. 10.1038/s41467-020-18560-z

Oksanen, J., Blanchet, F.G., Friendly, M., Kindt, R., Legendre, P., McGlinn, D., Minchin, P., O’Hara, R.B., Simpson, G., Solymos, P., Stevens, M.H.H., Szöcs, E., Wagner, H., 2020. vegan community ecology package version 2.5-7 November 2020.

Oktavia, D., Jin, G., 2020. Variations in leaf morphological and chemical traits in response to life stages, plant functional types, and habitat types in an old-growth temperate forest. Basic Appl. Ecol. 49, 22–33. 10.1016/j.baae.2020.09.010

Pantanella, F., Berlutti, F., Passariello, C., Sarli, S., Morea, C., Schippa, S., 2006. Violacein and biofilm production in Janthinobacterium lividum. J. Appl. Microbiol. 061120055200056-??? 10.1111/j.1365-2672.2006.03155.x

Patzak, R., Richter, R., Engelmann, R.A., Wirth, C., 2020. Tree crowns as meeting points of diversity generating mechanisms – a test with epiphytic lichens in a temperate forest. 10.1101/2020.01.03.894303

Ponette-González, A.G., Van Stan II, J.T., Magyar, D., 2020. Things Seen and Unseen in Throughfall and Stemflow, in: Precipitation Partitioning by Vegetation. Springer International Publishing, Cham, pp. 71–88. 10.1007/978-3-030-29702-2_5

Posfai, A., Taillefumier, T., Wingreen, N.S., 2017. Metabolic Trade-Offs Promote Diversity in a Model Ecosystem. Phys. Rev. Lett. 118, 028103. 10.1103/PhysRevLett.118.028103

Ren, Y., Xun, W., Yan, H., Ma, A., Xiong, W., Shen, Q., Zhang, R., 2020. Functional compensation dominates the assembly of plant rhizospheric bacterial community. Soil Biol. Biochem. 150, 107968. 10.1016/j.soilbio.2020.107968

Richter, R., Ballasus, H., Engelmann, R.A., Zielhofer, C., Sanaei, A., Wirth, C., 2022. Tree species matter for forest microclimate regulation during the drought year 2018: disentangling environmental drivers and biotic drivers. Sci. Rep. 12, 17559. 10.1038/s41598-022-22582-6

Richter, R., Reu, B., Wirth, C., Doktor, D., Vohland, M., 2016. The use of airborne hyperspectral data for tree species classification in a species-rich Central European forest area. Int. J. Appl. Earth Obs. Geoinf. 52, 464–474. 10.1016/j.jag.2016.07.018

Sancho-Knapik, D., Martín-Sánchez, R., Alonso-Forn, D., Peguero-Pina, J.J., Ferrio, J.P., Gil-Pelegrín, E., 2023. Trade-offs among leaf toughness, constitutive chemical defense, and growth rates in oaks are influenced by the level of leaf mass per area. Ann. For. Sci. 80, 39. 10.1186/s13595-023-01204-9

Schäfer, M., Vogel, C.M., Bortfeld-Miller, M., Mittelviefhaus, M., Vorholt, J.A., 2022. Mapping phyllosphere microbiota interactions in planta to establish genotype–phenotype relationships. Nat. Microbiol. 7, 856–867. 10.1038/s41564-022-01132-w

Schlechter, R.O., Kear, E.J., Bernach, M., Remus, D.M., Remus-Emsermann, M.N.P., 2023. Metabolic resource overlap impacts competition among phyllosphere bacteria. ISME J. 17, 1445–1454. 10.1038/s41396-023-01459-0

Stegen, J.C., Lin, X., Fredrickson, J.K., Chen, X., Kennedy, D.W., Murray, C.J., Rockhold, M.L., Konopka, A., 2013. Quantifying community assembly processes and identifying features that impose them. ISME J. 7, 2069–2079. 10.1038/ismej.2013.93

Stone, B.W.G., Jackson, C.R., 2019. Canopy position is a stronger determinant of bacterial community composition and diversity than environmental disturbance in the phyllosphere. FEMS Microbiol. Ecol. 95. 10.1093/femsec/fiz032

Su, L., Zhao, C., Xu, W., Xie, Z., 2019. Hydrochemical Fluxes in Bulk Precipitation, Throughfall, and Stemflow in a Mixed Evergreen and Deciduous Broadleaved Forest. Forests 10, 507. 10.3390/f10060507

Tamas, I., Smirnova, A. V, He, Z., Dunfield, P.F., 2014. The (d)evolution of methanotrophy in the Beijerinckiaceae —a comparative genomics analysis. ISME J. 8, 369–382. 10.1038/ismej.2013.145

Teachey, M., Pound, P., Ottesen, E., Van Stan, J., 2018. Bacterial Community Composition of Throughfall and Stemflow. Front. For. Glob. Chang. 1, 7. 10.3389/ffgc.2018.00007

Tukey, H.B., 1970. The Leaching of Substances from Plants. Annu. Rev. Plant Physiol. 21, 305–324. 10.1146/annurev.pp.21.060170.001513

Van Stan, J.T., Levia, D.F., Inamdar, S.P., Lepori-Bui, M., Mitchell, M.J., 2012. The effects of phenoseason and storm characteristics on throughfall solute washoff and leaching dynamics from a temperate deciduous forest canopy. Sci. Total Environ. 430, 48–58. 10.1016/j.scitotenv.2012.04.060

Venkatachalam, S., Ranjan, K., Prasanna, R., Ramakrishnan, B., Thapa, S., Kanchan, A., 2016. Diversity and functional traits of culturable microbiome members, including cyanobacteria in the rice phyllosphere. Plant Biol. 18, 627–637. 10.1111/plb.12441

Vermeuel, M.P., Novak, G.A., Kilgour, D.B., Claflin, M.S., Lerner, B.M., Trowbridge, A.M., Thom, J., Cleary, P.A., Desai, A.R., Bertram, T.H., 2023. Observations of biogenic volatile organic compounds over a mixed temperate forest during the summer to autumn transition. Atmos. Chem. Phys. 23, 4123–4148. 10.5194/acp-23-4123-2023

Wang, J., Shi, X., Lucas-Borja, M.E., Guo, Q., Wang, L., Huang, Z., 2023. Contribution of tree species to the co-occurrence network of the leaf phyllosphere and soil bacterial community in the subtropical forests. J. Environ. Manage. 343, 118274. 10.1016/j.jenvman.2023.118274

Wong, C.Y.S., D’Odorico, P., Arain, M.A., Ensminger, I., 2020. Tracking the phenology of photosynthesis using carotenoid-sensitive and near-infrared reflectance vegetation indices in a temperate evergreen and mixed deciduous forest. New Phytol. 226, 1682–1695. 10.1111/nph.16479

Woo, C., Yamamoto, N., 2020. Falling bacterial communities from the atmosphere. Environ. Microbiome 15, 22. 10.1186/s40793-020-00369-4

Yin, Y., Wang, Y.-F., Cui, H.-L., Zhou, R., Li, L., Duan, G.-L., Zhu, Y.-G., 2023. Distinctive Structure and Assembly of Phyllosphere Microbial Communities between Wild and Cultivated Rice. Microbiol. Spectr. 11, e0437122. 10.1128/spectrum.04371-22

Zhang, Y., Du, R., Chen, H., Du, P., Zhang, S., Ren, W., 2022. Different characteristics of microbial diversity and special functional microbes in rainwater and topsoil before and after 2019 new coronavirus epidemic in Inner Mongolia Grassland. Sci. Total Environ. 809, 151088. 10.1016/j.scitotenv.2021.151088

Zhou, J., Ning, D., 2017. Stochastic Community Assembly: Does It Matter in Microbial Ecology? Microbiol. Mol. Biol. Rev. 81. 10.1128/MMBR.00002-17

Zhou, S.-Y.-D., Li, H., Giles, M., Neilson, R., Yang, X., Su, J., 2021. Microbial Flow Within an Air-Phyllosphere-Soil Continuum. Front. Microbiol. 11. 10.3389/fmicb.2020.615481

