## Supplementary figures and tables for "Traversing the canopy: phenology-driven changes and within-canopy transport shape the phyllosphere microbiome in a temperate floodplain hardwood forest"

**Figure S1.** Shannon's diversity index H'. 1. across phenological stages (a, b and c), and 2. at different canopy positions and phenological stages (d, e and f) for the phyllosphere communities. The alphabets denote the statistical significance based on Kruskal–Wallis and Dunn's pairwise test, $P<0.05$.

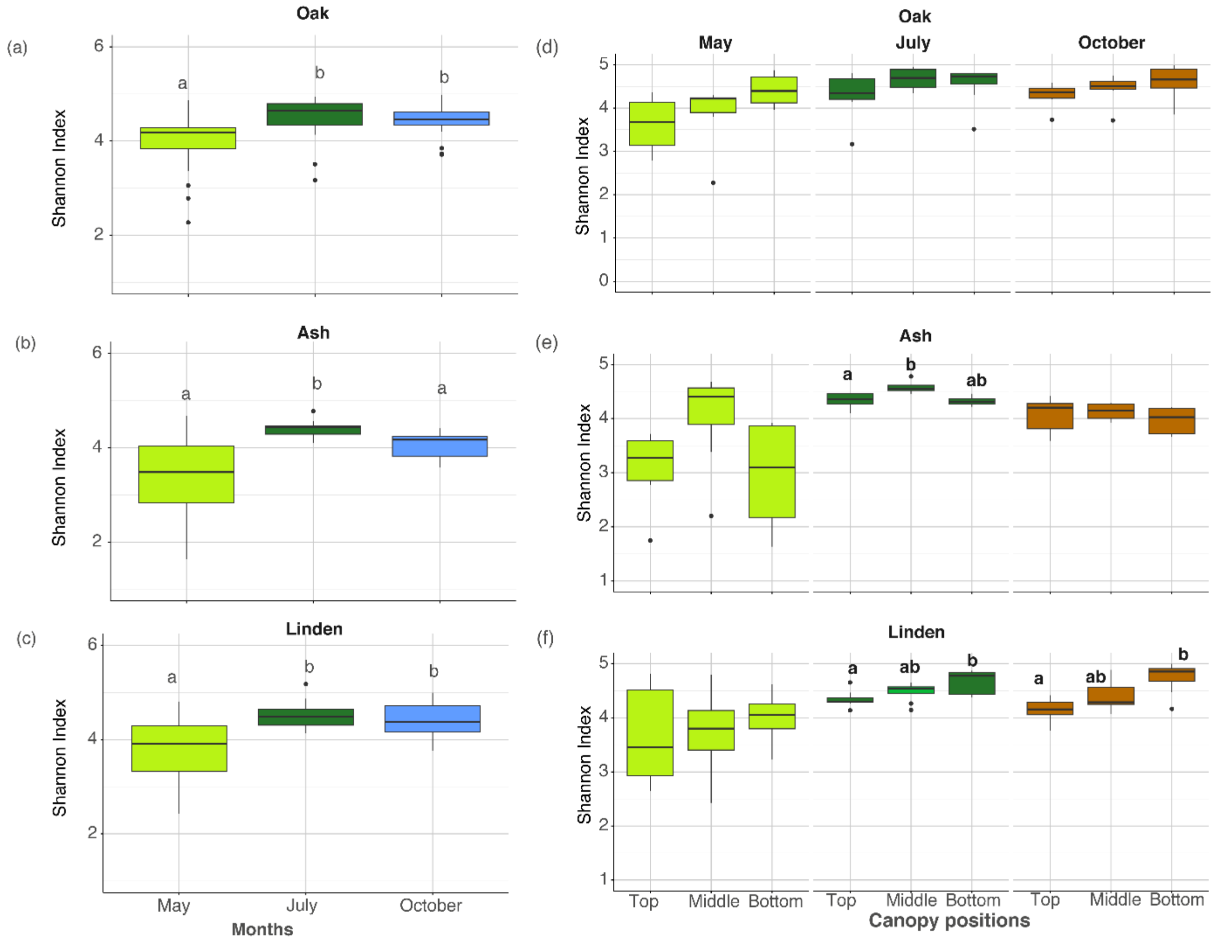

**Figure S2.** The relative abundance of bacterial families in the phyllosphere communities across canopy positions, phenological stages and tree species.

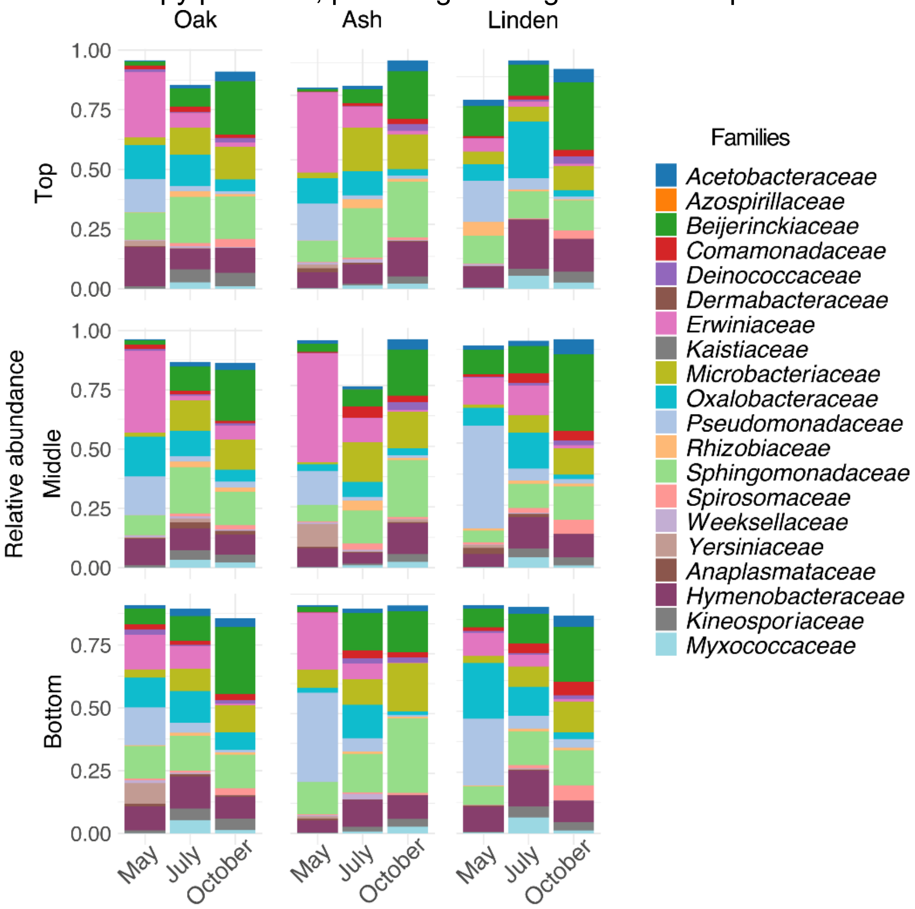

**Figure S3.** The estimated cell abundances of phyllosphere bacteria belonging to *Beijerinkiaceae* (a, b and c) and *Erwiniaceae* (d, e and f) families across tree species and canopy positions. Alphabets denote statistical differences between canopy positions. $^{\$}$ depicts Kruskal's wallis and Dunn's pairwise test ($P$<0.05).

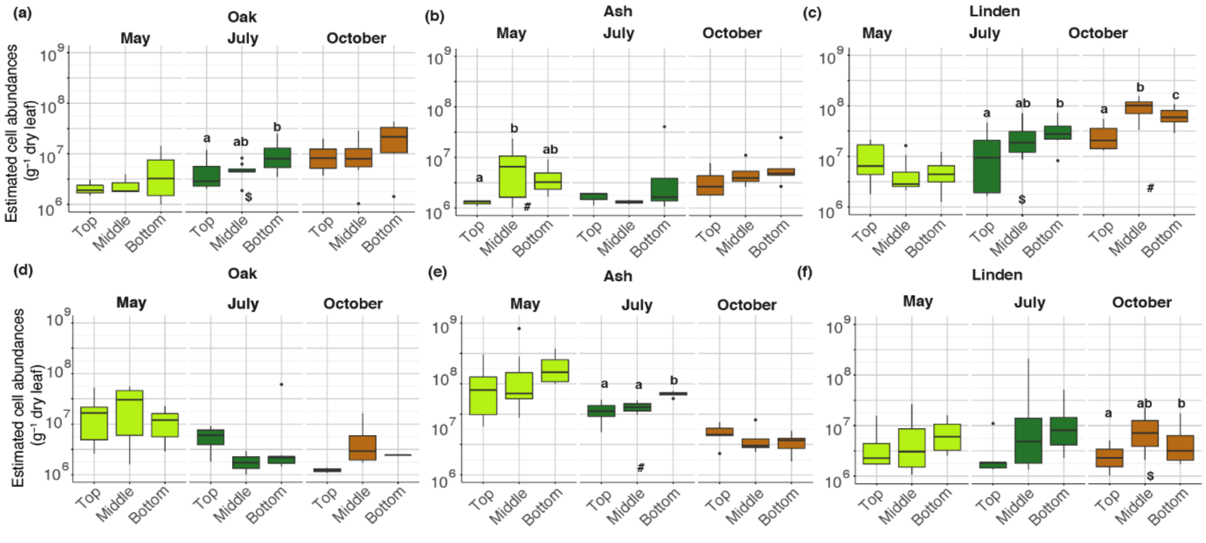

**Figure S4.** The relative percentage of successional and newly introduced ASVs across the phenological succession. July represents the May to July succession, while October represents the July to October succession.

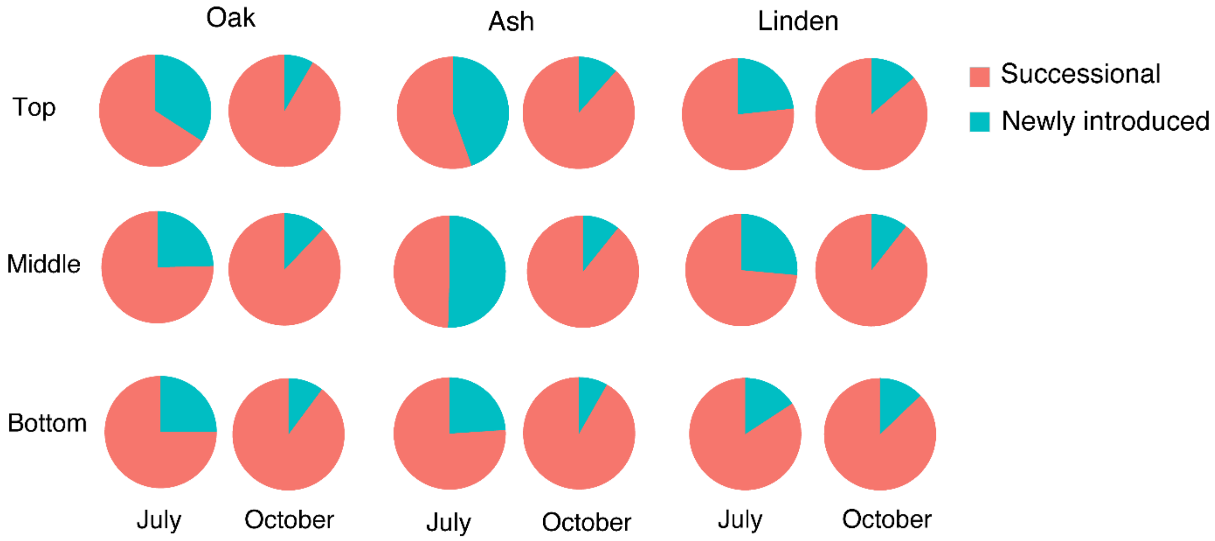

**Figure S5. Relative proportions of ASV categories (abundant, intermediate, rare) across phenological stages for (a) successional and (b) newly introduced ASVs.** July represents the May to July succession, while October represents the July to October succession.

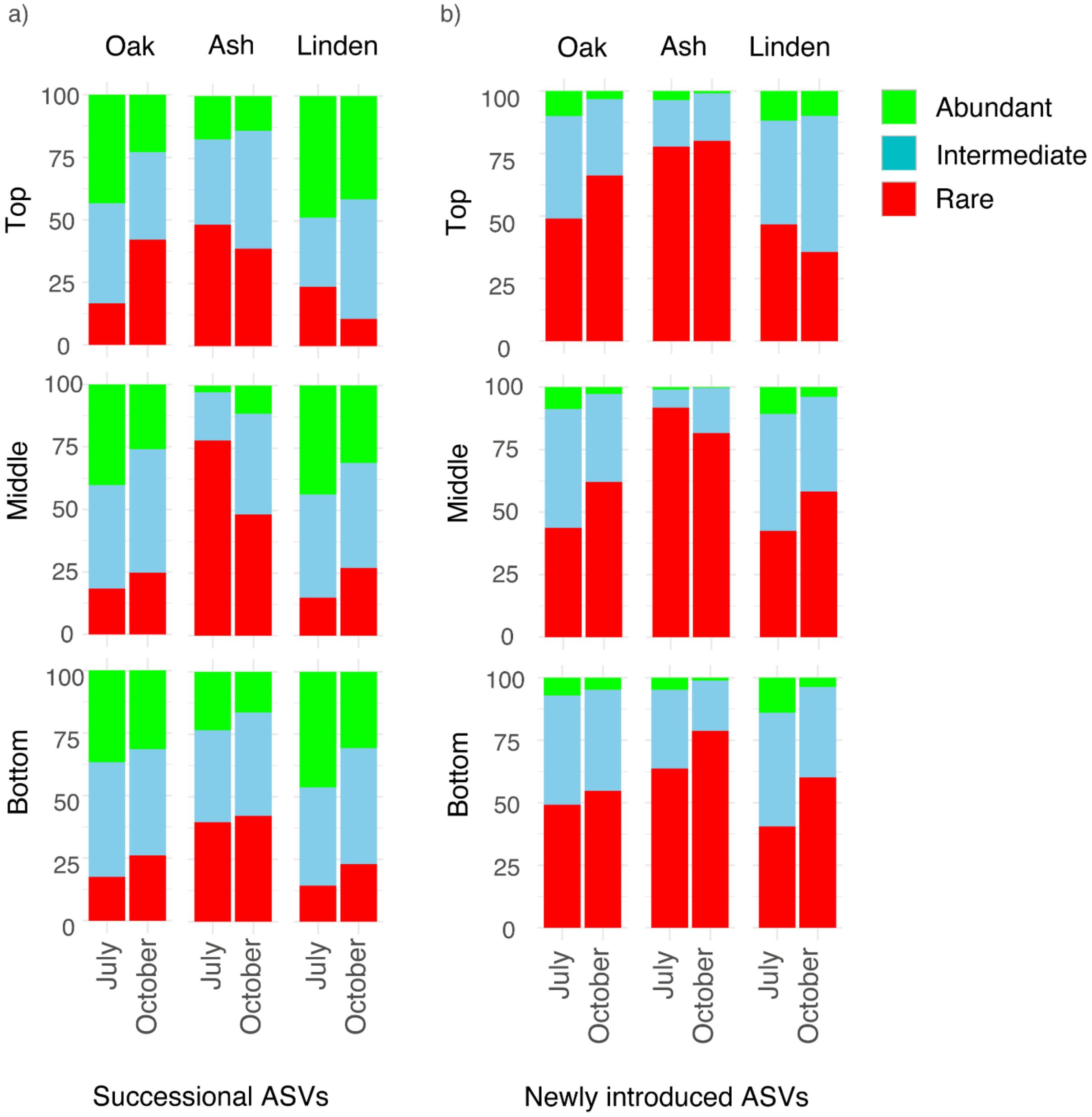

**Figure S6.** Shannon's diversity index H'. 1. across phenological stages (a, b and c), and 2. further resolved according to different canopy positions (d, e and f) for throughfall. The alphabets denote the statistical significance based on Kruskal–Wallis and Dunn's pairwise test, *P*<0.05.

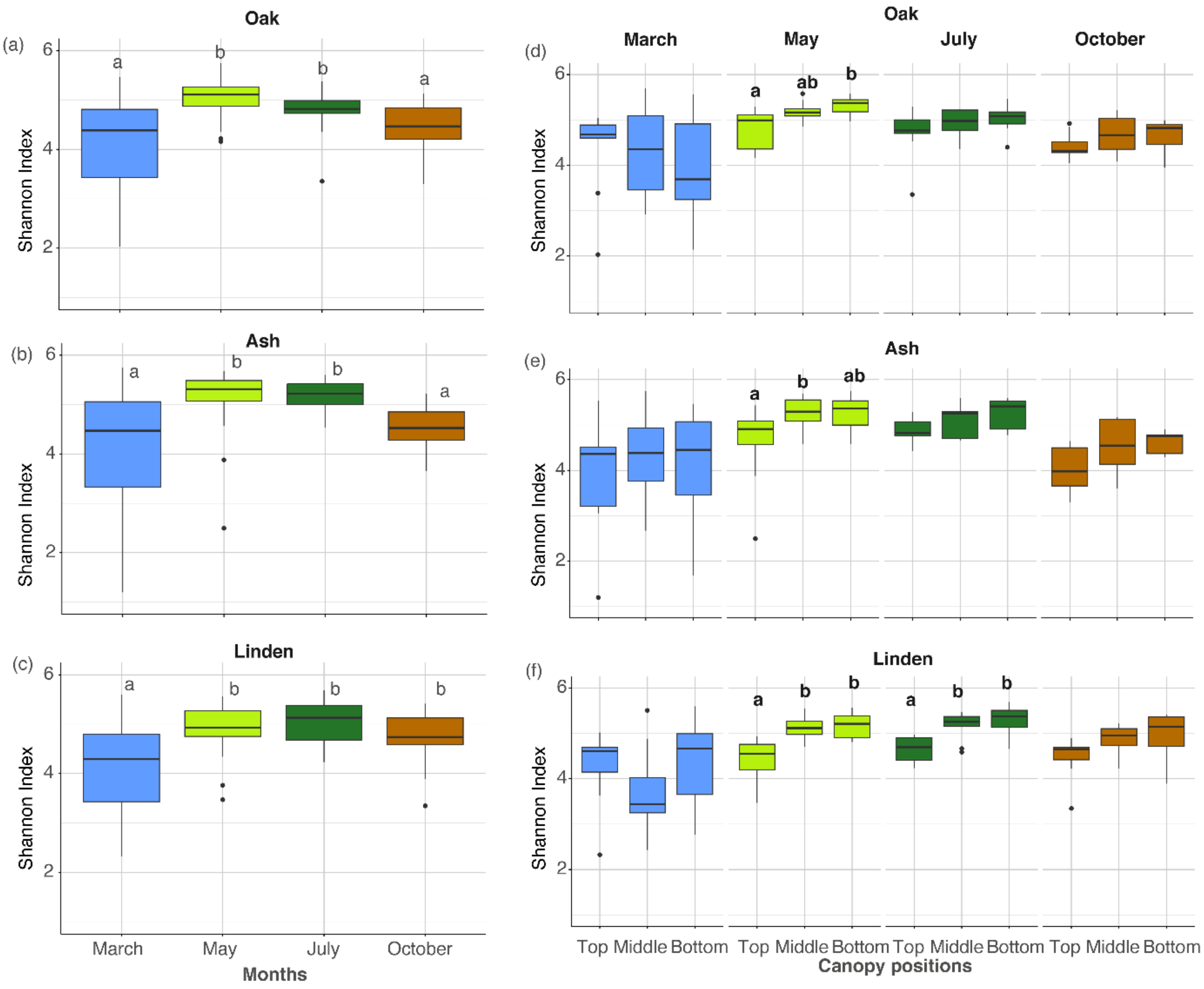

**Figure S7.** Throughfall bacterial flux across canopy positions, phenological stages and tree species. Alphabets denote statistical differences between positions. # indicates one-way ANOVA and Tukey's HSD pairwise test (*P*<0.05), while $ depicts statistical significance (*P*<0.05) of Kruskal's wallis and Dunn's pairwise test (*P*<0.05).

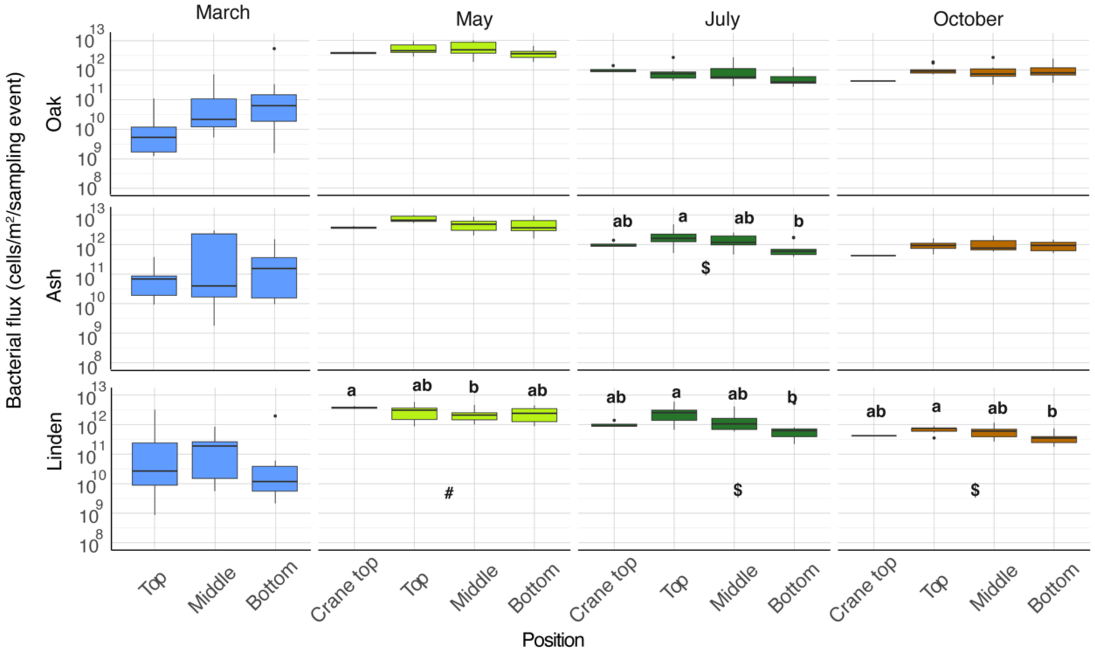

**Figure S8.** The relative abundance of bacterial families in throughfall across canopy positions, phenological stages and tree species.

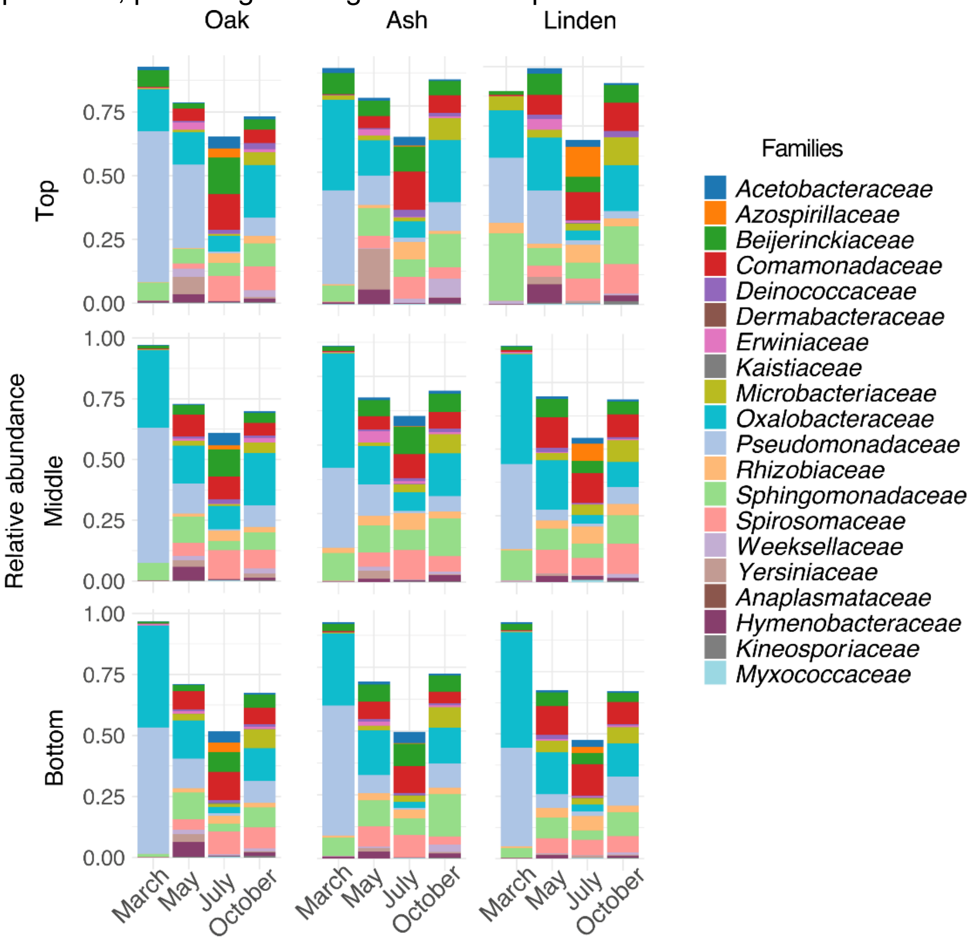

**Supplementary tables:**

**Table S1.** Metadata (Latitude, Longitude, Diameter at Breast Height (DBH), Altitude) of the tree individuals samples from Leipzig Canopy Crane facility, Germany.

| **Tree individual** | **Tree species** | **Latitude** | **Longitude** | **DBH (cm)** | **Altitude (m NN)** | **Predicted age (y)** | **Minimum age (y)** |
|---|---|---|---|---|---|---|---|
| ID129 | *Quercus robur* | 51.3651972 | 12.3089685 | 136.7 | 102 | 370 | - |
| ID131 | *Quercus robur* | 51.3651568 | 12.3088270 | 100.6 | 102 | - | - |
| ID733 | *Quercus robur* | 51.3651312 | 12.3098006 | 55.5 | 102 | 102 | - |
| ID232 | *Fraxinus excelsior* | 51.3656189 | 12.3090301 | 68.7 | 102 | - | - |
| ID435 | *Fraxinus excelsior* | 51.3663795 | 12.3092601 | 81.5 | 102 | - | - |
| ID770 | *Fraxinus excelsior* | 51.3650268 | 12.3095312 | 84.0 | 102 | - | - |
| ID439 | *Tilia cordata* | 51.3662962 | 12.3091490 | 76.8 | 102 | - | - |
| ID455 | *Tilia cordata* | 51.3663526 | 12.3096566 | 71 | 102 | 205 | 129 |
| ID517 | *Tilia cordata* | 51.3662235 | 12.3098308 | 81 | 102 | 235 | 148 |

**Table S2.** PERMANOVA analysis for the effects of Month, Host species, Position, Tree individuals and their interactions on bacterial communities of a) phyllosphere, b) throughfall bacterial communities c) month-wise for phyllosphere. $R^2$: proportion of variance explained by each factor. F: F-statistic indicating the strength of the effect. Pr(>F): p-value, with significance levels indicated as: *** p < 0.001, ** p < 0.01, * p < 0.05. Unexplained: proportion of variance not accounted for by the model.

a)

| | $R^2$ | F | Pr(>F) |
|---|---|---|---|
| Month | 19.236 | 30.5311 | 0.001 *** |
| Host | 12.18 | 19.3319 | 0.001 *** |
| Position | 2.343 | 3.7188 | 0.001 *** |
| Tree individual | 1.151 | 3.6544 | 0.001 *** |
| Month*host | 6.081 | 4.8258 | 0.001 *** |
| Host*Position | 1.792 | 1.4219 | 0.017 * |
| Month*Position | 1.772 | 1.4066 | 0.015 * |
| Unexplained | 55.444 | | |

b)

| | $R^2$ | F | Pr(>F) |
|---|---|---|---|
| Month | 19.984 | 31.7697 | 0.001*** |
| Host | 4.384 | 10.4536 | 0.001*** |
| Position | 2.221 | 5.2955 | 0.001*** |
| Tree individual | 3.896 | 3.0972 | 0.001*** |
| Month*host | 4.496 | 3.574 | 0.001*** |
| Host*Position | 2.944 | 2.3401 | 0.001*** |
| Month*Position | 1.479 | 1.7633 | 0.001*** |
| Unexplained | 60.596 | | |

c)

| | $R^2$ | F | Pr(>F) |
|---|---|---|---|
| **May** | | | |
| Host | 19.48 | 7.5996 | 0.001 *** |
| Position | 4.9 | 1.9308 | 0.001 *** |
| Tree individual | 2.51 | 1.965 | 0.007 ** |
| **July** | | | |
| Host | 23.66 | 10.6557 | 0.001 *** |
| Position | 5.86 | 2.6393 | 0.001 *** |
| Tree individual | 1.48 | 1.4811 | 0.098 . |
| **October** | | | |
| Host | 28.37 | 11.74 | 0.001 *** |
| Position | 5 | 2.0714 | 0.004 ** |

| | | | |
|---|---|---|---|
| Tree individual | 4.9 | 4.1054 | 0.001 *** |

**Table S3.** The statistical tests for estimated bacterial cell abundances of a) phyllosphere, b) throughfall, c) *Beijerinkiaceae* and d) *Erwiniaceae* families across tree species and canopy positions. Alphabets denote statistical difference between canopy positions based on Kruskal's wallis and Dunn's pairwise test ($P$<0.05). $^{\$}$ indicates one-way ANOVA and Tukey's HSD pairwise test ($P$<0.05), while $^{\#}$ depicts statistical significance ($P$<0.05) of Kruskal's wallis and Dunn's pairwise test ($P$<0.05). S indicates a significant while NS point to non-significant statistical test result.

a)

| Oak | Month | Position | Normality | Univariate test | Pairwise test |
|---|---|---|---|---|---|
| | May | Top | P<0.05 | NS | |
| | May | Middle | | | |
| | May | Bottom | | | |
| | July | Top | P>0.05 | S# | a |
| | July | Middle | | | ab |
| | July | Bottom | | | b |
| | October | Top | P<0.05 | NS | |
| | October | Middle | | | |
| | October | Bottom | | | |

| Ash | Month | Position | Normality | Univariate test | Pairwise test |
|---|---|---|---|---|---|
| | May | Top | P>0.05 | NS | |
| | May | Middle | | | |
| | May | Bottom | | | |
| | July | Top | P>0.05 | NS | |
| | July | Middle | | | |
| | July | Bottom | | | |
| | October | Top | P>0.05 | NS | |
| | October | Middle | | | |
| | October | Bottom | | | |

| Linden | Month | Position | Normality | Univariate test | Pairwise test |
|---|---|---|---|---|---|
| | May | Top | P<0.05 | NS | |
| | May | Middle | | | |
| | May | Bottom | | | |
| | July | Top | P<0.05 | NS | |
| | July | Middle | | | |
| | July | Bottom | | | |
| | October | Top | P>0.05 | S# | a |
| | October | Middle | | | b |
| | October | Bottom | | | b |

b)

| Oak | Month | Position | Normality | Univariate test | Pairwise test |
|---|---|---|---|---|---|
| | March | Top | P>0.05 | S# | a |
| | March | Middle | | | ab |
| | March | Bottom | | | b |
| | May | Top | P>0.05 | S# | NS |
| | May | Middle | | | |
| | May | Bottom | | | |
| | July | Top | P<0.05 | NS | |
| | July | Middle | | | |
| | July | Bottom | | | |
| | October | Top | P<0.05 | NS | |
| | October | Middle | | | |
| | October | Bottom | | | |

| Ash | Month | Position | Normality | Univariate test | Pairwise test |
|---|---|---|---|---|---|
| | March | Top | P>0.05 | S# | NS |
| | March | Middle | | | |
| | March | Bottom | | | |
| | May | Top | P<0.05 | NS | |
| | May | Middle | | | |
| | May | Bottom | | | |
| | July | Top | P<0.05 | NS | |
| | July | Middle | | | |
| | July | Bottom | | | |
| | October | Top | P>0.05 | S# | NS |
| | October | Middle | | | |
| | October | Bottom | | | |

| Linden | Month | Position | Normality | Univariate test | Pairwise test |
|---|---|---|---|---|---|
| | March | Top | P>0.05 | S# | NS |
| | March | Middle | | | |
| | March | Bottom | | | |
| | May | Top | P>0.05 | S# | NS |
| | May | Middle | | | |
| | May | Bottom | | | |
| | July | Top | P<0.05 | NS | |
| | July | Middle | | | |
| | July | Bottom | | | |
| | October | Top | P>0.05 | S# | NS |
| | October | Middle | | | |
| | October | Bottom | | | |

c)

| Oak | Month | Position | Normality | Univariate test | Pairwise test |
|---|---|---|---|---|---|
| | May | Top | P<0.05 | NS | |

| | | | | | |
|---|---|---|---|---|---|
| | May | Middle | | | |
| | May | Bottom | | | |
| | July | Top | P>0.05 | S# | a |
| | July | Middle | | | ab |
| | July | Bottom | | | b |
| | October | Top | P<0.05 | NS | |
| | October | Middle | | | |
| | October | Bottom | | | |

| Ash | Month | Position | Normality | Univariate test | Pairwise test |
|---|---|---|---|---|---|
| | May | Top | P<0.05 | S$ | a |
| | May | Middle | | | b |
| | May | Bottom | | | ab |
| | July | Top | P>0.05 | S# | NS |
| | July | Middle | | | |
| | July | Bottom | | | |
| | October | Top | P>0.05 | S# | NS |
| | October | Middle | | | |
| | October | Bottom | | | |

| Linden | Month | Position | Normality | Univariate test | Pairwise test |
|---|---|---|---|---|---|
| | May | Top | P>0.05 | S# | NS |
| | May | Middle | | | |
| | May | Bottom | | | |
| | July | Top | P>0.05 | S# | a |
| | July | Middle | | | ab |
| | July | Bottom | | | b |
| | October | Top | P<0.05 | S$ | a |
| | October | Middle | | | b |
| | October | Bottom | | | c |

d) Erwiniaceae

| Oak | Month | Position | Normality | Univariate test | Pairwise test |
|---|---|---|---|---|---|
| | May | Top | P<0.05 | NS | |
| | May | Middle | | | |
| | May | Bottom | | | |
| | July | Top | P<0.05 | NS | |
| | July | Middle | | | |
| | July | Bottom | | | |
| | October | Top | P>0.05 | S# | NS |
| | October | Middle | | | |
| | October | Bottom | | | |

| Ash | Month | Position | Normality | Univariate test | Pairwise test |
|---|---|---|---|---|---|
| | May | Top | P>0.05 | S | NS |
| | May | Middle | | | |
| | May | Bottom | | | |

| | | | | | |
|---|---|---|---|---|---|
| | July | Top | P<0.05 | S$ | a |
| | July | Middle | | | a |
| | July | Bottom | | | b |
| | October | Top | P>0.05 | S | NS |
| | October | Middle | | | |
| | October | Bottom | | | |

| Linden | Month | Position | Normality | Univariate test | Pairwise test |
|---|---|---|---|---|---|
| | May | Top | P>0.05 | S# | NS |
| | May | Middle | | | |
| | May | Bottom | | | |
| | July | Top | P>0.05 | S# | NS |
| | July | Middle | | | |
| | July | Bottom | | | |
| | October | Top | P>0.05 | S# | a |
| | October | Middle | | | ab |
| | October | Bottom | | | b |

**Table S4.** a) The relative percentage of successional and newly introduced ASVs across the phenological succession. July represents the May to July succession, while October represents the July to October succession. b) Relative proportions of ASV categories (abundant, intermediate, rare) across phenological stages for (a) successional and (b) newly introduced ASVs.

a)

| | Successional | New intro |
|---|---|---|
| Oak_Top_July | 65.8768549 | 34.1231451 |
| Oak_Top_October | 91.6227377 | 8.3772623 |
| Oak_Middle_July | 75.3740408 | 24.6259592 |
| Oak_Middle_October | 87.9001558 | 12.0998442 |
| Oak_Bottom_July | 75.0284232 | 24.9715768 |
| Oak_Bottom_October | 89.830558 | 10.169442 |
| Ash_Top_July | 55.4383747 | 44.5616253 |
| Ash_Top_October | 88.4132089 | 11.5867911 |
| Ash_Middle_July | 49.5445635 | 50.4554365 |
| Ash_Middle_October | 89.2402539 | 10.7597461 |
| Ash_Bottom_July | 76.0323371 | 23.9676629 |
| Ash_Bottom_October | 91.7582172 | 8.24178277 |
| Linden_Top_July | 76.690868 | 23.309132 |
| Linden_Top_October | 86.3617979 | 13.6382021 |
| Linden_Middle_July_ | 73.5716038 | 26.4283962 |
| Linden_Middle_October | 89.3617634 | 10.6382366 |
| Linden_Bottom_July | 84.1966843 | 15.8033157 |
| Linden_Bottom_October | 87.1918877 | 12.8081123 |

b)

| Successional ASV | Abundant | Intermediate | Rare |
|---|---|---|---|
| Oak_Top_July | 43.3823529 | 40.0735294 | 16.5441176 |
| Oak_Top_October | 22.9591837 | 35.0340136 | 42.0068027 |
| Oak_Middle_July | 40.2857143 | 41.4285714 | 18.2857143 |
| Oak_Middle_October | 25.9504132 | 49.4214876 | 24.6280992 |
| Oak_Bottom_July | 36.6754617 | 45.9102902 | 17.414248 |
| Oak_Bottom_October | 31.5068493 | 42.4657534 | 26.0273973 |
| Ash_Top_July | 17.3267327 | 34.1584158 | 48.5148515 |
| Ash_Top_October | 13.9240506 | 47.1518987 | 38.9240506 |
| Ash_Middle_July | 2.74509804 | 19.2156863 | 78.0392157 |

| | | | |
|---|---|---|---|
| Ash_Middle_October | 11.2781955 | 40.2255639 | 48.4962406 |
| Ash_Bottom_July | 23.3830846 | 36.8159204 | 39.800995 |
| Ash_Bottom_October | 16.2711864 | 41.3559322 | 42.3728814 |
| Linden_Top_July | 48.6607143 | 27.6785714 | 23.6607143 |
| Linden_Top_October | 41.260745 | 47.8510029 | 10.8882521 |
| Linden_Middle_July | 43.5810811 | 41.2162162 | 15.2027027 |
| Linden_Middle_October | 30.9090909 | 42.0454545 | 27.0454545 |
| Linden_Bottom_July | 46.2462462 | 39.3393393 | 14.4144144 |
| Linden_Bottom_October | 30.5376344 | 46.4516129 | 23.0107527 |

| Newly introduced ASVs | Abundant | Intermediate | Rare |
|---|---|---|---|
| Oak_Top_July | 10.0470958 | 40.9733124 | 48.9795918 |
| Oak_Top_October | 3.24675325 | 30.5194805 | 66.2337662 |
| Oak_Middle_July | 8.77513711 | 47.5319927 | 43.6928702 |
| Oak_Middle_October | 2.81124498 | 35.1405622 | 62.0481928 |
| Oak_Bottom_July | 7.07803993 | 43.738657 | 49.1833031 |
| Oak_Bottom_October | 4.81400438 | 40.4814004 | 54.7045952 |
| Ash_Top_July | 3.68906456 | 18.5770751 | 77.7338603 |
| Ash_Top_October | 0.93896714 | 19.0140845 | 80.0469484 |
| Ash_Middle_July | 0.99573257 | 7.25462304 | 91.7496444 |
| Ash_Middle_October | 0.33444816 | 18.0602007 | 81.6053512 |
| Ash_Bottom_July | 4.83870968 | 31.4516129 | 63.7096774 |
| Ash_Bottom_October | 1.179941 | 20.0589971 | 78.7610619 |
| Linden_Top_July | 11.8729097 | 41.4715719 | 46.6555184 |
| Linden_Top_October | 9.91957105 | 54.4235925 | 35.6568365 |
| Linden_Middle_July | 10.7655502 | 46.7703349 | 42.4641148 |
| Linden_Middle_October | 3.88548057 | 37.8323108 | 58.2822086 |
| Linden_Bottom_July | 13.9874739 | 45.5114823 | 40.5010438 |
| Linden_Bottom_October | 3.73514431 | 36.1629881 | 60.1018676 |

**Table S5.** Amplicon sequence variants (ASVs) detected only during a) July or b) October at tree species and canopy position levels

a)

| Host | ASV | Family | Genus |
|---|---|---|---|
| Oak_Top | ASV89 | Microbacteriaceae | Rathayibacter |
| Oak_Top | ASV47 | Nocardiaceae | Rhodococcus |
| Oak_Top | ASV50 | Sphingobacteriaceae | Pedobacter |
| Oak_Top | ASV45 | Oxalobacteraceae | Massilia |
| Oak_Middle | ASV47 | Nocardiaceae | Rhodococcus |
| Oak_Middle | ASV49 | Sphingobacteriaceae | Pedobacter |
| Oak_Middle | ASV25 | Sphingobacteriaceae | Pedobacter |
| Oak_Bottom | ASV50 | Sphingobacteriaceae | Pedobacter |
| Ash_Top | ASV89 | Microbacteriaceae | Rathayibacter |
| Ash_Top | ASV47 | Nocardiaceae | Rhodococcus |
| Ash_Top | ASV49 | Sphingobacteriaceae | Pedobacter |
| Ash_Top | ASV94 | Oxalobacteraceae | Massilia |
| Ash_Top | ASV87 | Comamonadaceae | Variovorax |
| Ash_Middle | ASV71 | Rhizobiaceae | Allorhizobium-Neorhizobium-Pararhizobium-Rhizobium |
| Ash_Middle | ASV89 | Microbacteriaceae | Rathayibacter |
| Ash_Middle | ASV47 | Nocardiaceae | Rhodococcus |
| Ash_Middle | ASV40 | Rhizobiaceae | Ensifer |
| Ash_Middle | ASV28 | Comamonadaceae | Variovorax |
| Ash_Middle | ASV95 | Beijerinckiaceae | |
| Ash_Middle | ASV84 | Spirosomaceae | Spirosoma |
| Ash_Middle | ASV46 | Xanthobacteraceae | Rhodopseudomonas |
| Ash_Bottom | ASV81 | Morganellaceae | Buchnera |
| Ash_Bottom | ASV52 | Oxalobacteraceae | Massilia |
| Ash_Bottom | ASV47 | Nocardiaceae | Rhodococcus |
| Ash_Bottom | ASV65 | Oxalobacteraceae | Massilia |
| Ash_Bottom | ASV87 | Comamonadaceae | Variovorax |
| Linden_Top | ASV77 | Hymenobacteraceae | Hymenobacter |
| Linden_Top | ASV100 | Oxalobacteraceae | Massilia |
| Linden_Top | ASV42 | Microbacteriaceae | |
| Linden_Top | ASV89 | Microbacteriaceae | Rathayibacter |

| | | | |
|---|---|---|---|
| Linden_Top | ASV96 | Yersiniaceae | Serratia |
| Linden_Top | ASV33 | Yersiniaceae | Serratia |
| Linden_Top | ASV39 | Oxalobacteraceae | Massilia |
| Linden_Middle | ASV81 | Morganellaceae | Buchnera |
| Linden_Middle | ASV90 | Oxalobacteraceae | Massilia |
| Linden_Middle | ASV100 | Oxalobacteraceae | Massilia |
| Linden_Middle | ASV40 | Rhizobiaceae | Ensifer |
| Linden_Middle | ASV62 | Oxalobacteraceae | Massilia |
| Linden_Middle | ASV28 | Comamonadaceae | Variovorax |
| Linden_Middle | ASV84 | Spirosomaceae | Spirosoma |
| Linden_Middle | ASV46 | Xanthobacteraceae | Rhodopseudomonas |
| Linden_Middle | ASV89 | Microbacteriaceae | Rathayibacter |
| Linden_Middle | ASV39 | Oxalobacteraceae | Massilia |
| Linden_Middle | ASV72 | Oxalobacteraceae | Duganella |
| Linden_Middle | ASV60 | Pseudomonadaceae | Pseudomonas |
| Linden_Middle | ASV59 | Oxalobacteraceae | Janthinobacterium |
| Linden_Middle | ASV75 | Spirosomaceae | Spirosoma |
| Linden_Middle | ASV78 | Hymenobacteraceae | Hymenobacter |
| Linden_Bottom | ASV70 | Erwiniaceae | Pantoea |
| Linden_Bottom | ASV47 | Nocardiaceae | Rhodococcus |
| Linden_Bottom | ASV100 | Oxalobacteraceae | Massilia |
| Linden_Bottom | ASV89 | Microbacteriaceae | Rathayibacter |
| Linden_Bottom | ASV83 | Sphingobacteriaceae | Pedobacter |
| Linden_Bottom | ASV75 | Spirosomaceae | Spirosoma |

b)

| Host | ASV | Family | Genus |
|---|---|---|---|
| Oak_Top | ASV179 | Beijerinckiaceae | 1174-901-12 |
| Oak_Top | ASV125 | Acetobacteraceae | Acidiphilium |
| Oak_Top | ASV69 | Pseudomonadaceae | Pseudomonas |
| Oak_Top | ASV145 | Pseudomonadaceae | Pseudomonas |
| Oak_Top | ASV92 | Acetobacteraceae | |
| Oak_Top | ASV141 | Spirosomaceae | Spirosoma |
| Oak_Middle | ASV27 | Erwiniaceae | Erwinia |
| Oak_Middle | ASV145 | Pseudomonadaceae | Pseudomonas |
| Oak_Bottom | ASV151 | Pseudomonadaceae | Pseudomonas |
| Oak_Bottom | ASV55 | Erwiniaceae | Erwinia |
| Oak_Bottom | ASV135 | Oxalobacteraceae | Massilia |
| Oak_Bottom | ASV182 | Spirosomaceae | Spirosoma |
| Oak_Bottom | ASV41 | Pseudomonadaceae | Pseudomonas |
| Ash_Top | ASV102 | Acetobacteraceae | Acidiphilium |

| | | | |
|---|---|---|---|
| Ash_Top | ASV178 | Spirosomaceae | Spirosoma |
| Ash_Top | ASV99 | Beijerinckiaceae | Methylocella |
| Ash_Top | ASV179 | Beijerinckiaceae | 1174-901-12 |
| Ash_Middle | ASV164 | Myxococcaceae | P3OB-42 |
| Ash_Middle | ASV150 | Spirosomaceae | Huanghella |
| Ash_Middle | ASV137 | Hymenobacteraceae | Hymenobacter |
| Ash_Middle | ASV176 | Acetobacteraceae | Acidiphilium |
| Ash_Middle | ASV7 | Oxalobacteraceae | Massilia |
| Ash_Bottom | ASV131 | Sphingomonadaceae | Sphingomonas |
| Ash_Bottom | ASV178 | Spirosomaceae | Spirosoma |
| Linden_Top | ASV134 | Morganellaceae | Incertae Sedis |
| Linden_Top | ASV193 | Sphingomonadaceae | Sphingomonas |
| Linden_Top | ASV55 | Erwiniaceae | Erwinia |
| Linden_Top | ASV141 | Spirosomaceae | Spirosoma |
| Linden_Top | ASV83 | Sphingobacteriaceae | Pedobacter |
| Linden_Middle | ASV134 | Morganellaceae | Incertae Sedis |
| Linden_Middle | ASV83 | Sphingobacteriaceae | Pedobacter |
| Linden_Bottom | ASV182 | Spirosomaceae | Spirosoma |
| Linden_Bottom | ASV121 | Erwiniaceae | Izhakiella |
| Linden_Bottom | ASV125 | Acetobacteraceae | Acidiphilium |
| Linden_Bottom | ASV10 | Pseudomonadaceae | Pseudomonas |
| Linden_Bottom | ASV134 | Morganellaceae | Incertae Sedis |
| Linden_Bottom | ASV102 | Acetobacteraceae | Acidiphilium |
| Linden_Bottom | ASV189 | Spirosomaceae | Spirosoma |

**Table S6.** Throughfall bacterial flux across canopy positions, phenological stages and tree species. Alphabets denote statistical differences between positions. # indicates one-way ANOVA and Tukey's HSD pairwise test (*P*<0.05), while $ depicts statistical significance (*P*<0.05) of Kruskal's wallis and Dunn's pairwise test (*P*<0.05). S indicates a significant while NS point to non-significant statistical test result.

| Oak | Month | Position | Normality | Univariate test | Pairwise-Test |
|---|---|---|---|---|---|
| | March | Top | P>0.05 | S $ | NS |
| | March | Middle | | | |
| | March | Bottom | | | |
| | May | Crane top | P>0.05 | S $ | NS |
| | May | Top | | | |
| | May | Middle | | | |
| | May | Bottom | | | |
| | July | Crane top | P>0.05 | S $ | NS |
| | July | Top | | | |
| | July | Middle | | | |
| | July | Bottom | | | |
| | October | Crane top | P>0.05 | S $ | NS |
| | October | Top | | | |
| | October | Middle | | | |
| | October | Bottom | | | |

| Ash | Month | Position | Normality | Univariate test | Pairwise-Test |
|---|---|---|---|---|---|
| | March | Top | P>0.05 | S $ | NS |
| | March | Middle | | | |
| | March | Bottom | | | |
| | May | Crane top | P>0.05 | S $ | NS |
| | May | Top | | | |
| | May | Middle | | | |
| | May | Bottom | | | |
| | July | Crane top | P>0.05 | S $ | ab |
| | July | Top | | | a |
| | July | Middle | | | ab |
| | July | Bottom | | | b |
| | October | Crane top | P>0.05 | S $ | NS |
| | October | Top | | | |
| | October | Middle | | | |
| | October | Bottom | | | |

| Linden | Month | Position | Normality | Univariate test | Pairwise-Test |
|---|---|---|---|---|---|
| | March | Top | P>0.05 | S $ | NS |

| | | | | | |
|---|---|---|---|---|---|
| | March | Middle | | | |
| | March | Bottom | | | |
| | May | Crane top | $P<0.05$ | S # | a |
| | May | Top | | | ab |
| | May | Middle | | | b |
| | May | Bottom | | | ab |
| | July | Crane top | $P>0.05$ | S $ | ab |
| | July | Top | | | a |
| | July | Middle | | | ab |
| | July | Bottom | | | b |
| | October | Crane top | $P>0.05$ | S$ | ab |
| | October | Top | | | a |
| | October | Middle | | | ab |
| | October | Bottom | | | b |
